# Lineage tracing of human embryonic development and foetal haematopoiesis through somatic mutations

**DOI:** 10.1101/2020.05.29.088765

**Authors:** Michael Spencer Chapman, Anna Maria Ranzoni, Brynelle Myers, Nick Williams, Tim Coorens, Emily Mitchell, Timothy Butler, Kevin J Dawson, Yvette Hooks, Luiza Moore, Jyoti Nangalia, Philip S Robinson, Elizabeth Hook, Peter J Campbell, Ana Cvejic

**Affiliations:** Wellcome Trust Sanger Institute, Wellcome Trust Genome Campus, Hinxton, CB10 1SA, UK; Department of Haematology, Hammersmith Hospital, Imperial College Healthcare NHS Trust, London, W12 0HS, UK; Department of Haematology, Cambridge University Hospitals NHS Foundation Trust, Cambridge, CB2 0QQ, UK; Wellcome Trust – Medical Research Council Cambridge Stem Cell Institute, Cambridge, CB2 0AW, UK; University of Cambridge, Department of Haematology, Cambridge, CB2 0AW, UK; Department of Histopathology, Cambridge University Hospitals NHS Foundation Trust, Cambridge, CB2 0QQ, UK; University of Cambridge, Department of Paediatrics, Cambridge, CB2 0QQ, UK

## Abstract

To date, ontogeny of the human haematopoietic system during foetal development has been characterized mainly through careful microscopic observations. Here we used whole-genome sequencing (WGS) of 511 single-cell derived haematopoietic colonies from healthy human foetuses of 8 and 18 post-conception weeks (pcw) coupled with deep targeted sequencing of tissues of known embryonic origin to reconstruct a phylogenetic tree of blood development. We found that in healthy foetuses, individual haematopoietic progenitors acquire tens of somatic mutations by 18 pcw. Using these mutations as barcodes, we timed the divergence of embryonic and extra-embryonic tissues during development and estimated the number of blood antecedents at different stages of embryonic development. Our analysis has shown that ectoderm originates from a smaller set of blood antecedents compared to endoderm and mesoderm. Finally, our data support a hypoblast origin of the extra-embryonic mesoderm and primitive blood in humans.

## Introduction

Human embryonic development starts with the division of the zygote to two daughter cells called blastomeres. After several further divisions, at approximately the 100-cell-stage, the cells physically separate into the inner cell mass (ICM) which is fated to become the embryo, and the trophoblast which develops into the chorionic sac and foetal component of the placenta. The ICM then undergoes further differentiation into the epiblast and hypoblast, which primarily contribute to embryonic and extra-embryonic tissues respectively. At 3 weeks of development, the epiblast differentiates into the three germ layers (ectoderm, mesoderm and endoderm). Ectoderm gives rise to the epidermis and nervous system; mesoderm generates the heart, kidney, skeleton, muscles and connective tissues; while endoderm forms the liver, lungs and inner layer of the digestive tract^1^.

The first haematopoietic cells (mainly nucleated erythrocytes and, to a lesser extent, macrophages and megakaryocytes^2–4^) start to form within blood islands in the yolk sac from 16 post-conception days (pcd). However, the developmental origin of these so-called primitive haematopoietic cells in humans is still debated. Yolk sac blood islands arise from the extra-embryonic mesoderm^5^. The precise origin of this extra-embryonic tissue remains unclear with some evidence in mice pointing to the primitive streak^6,7^, while histological studies in humans suggest it derives from the hypoblast^7,8^. Interestingly, human foetal histological studies have shown that at 8 pcw more than 90% of erythroid cells in circulating blood are still of primitive origin, while more than 80% of those found in liver derive from the aorta-gonad-mesonephros (the so-called ‘definitive’ wave of haematopoiesis). Thus, it has been speculated that definitive erythrocytes are temporarily retained in the liver^9^.

Establishment of the bone marrow, the main site of adult blood production, marks the end of the embryonic period in humans^4^. Haematopoietic colonisation of the bone marrow starts at ∼10.5 weeks with the appearance of monocytes/macrophages, followed by haematopoietic stem and progenitor cells (HSPCs), although the exact onset of haematopoietic stem cell (HSC) activity in the bone marrow and the kinetics of migration between different organs remain unclear^4,9,10^. Here, we used the accumulation of somatic mutations in foetal HSPCs to gain further insights into the dynamics of human prenatal development and the origins of primitive and definitive haematopoiesis.

### Reconstructing the phylogeny of foetal HSPCs from whole-genome sequencing data

Somatic mutations accumulate in a cell’s genome over successive cell divisions^11^. These mutations are inherited by all future progeny of the cell as they are copied during DNA replication. Therefore, somatic mutations can effectively be used as barcodes in lineage tracing studies^12^. By analysing patterns of mutations that have accumulated in somatic cells over time, it is possible to identify clonal relationships between cells and thereby reconstruct the phylogeny of a population^13^. In this study, we performed whole-genome sequencing (WGS) of single-cell-derived blood colonies from human foetal HSPCs, coupled with deep targeted sequencing of matched tissues of known embryonic origin, to build a phylogenetic tree of foetal development (Fig. 1a).

**Fig. 1.**
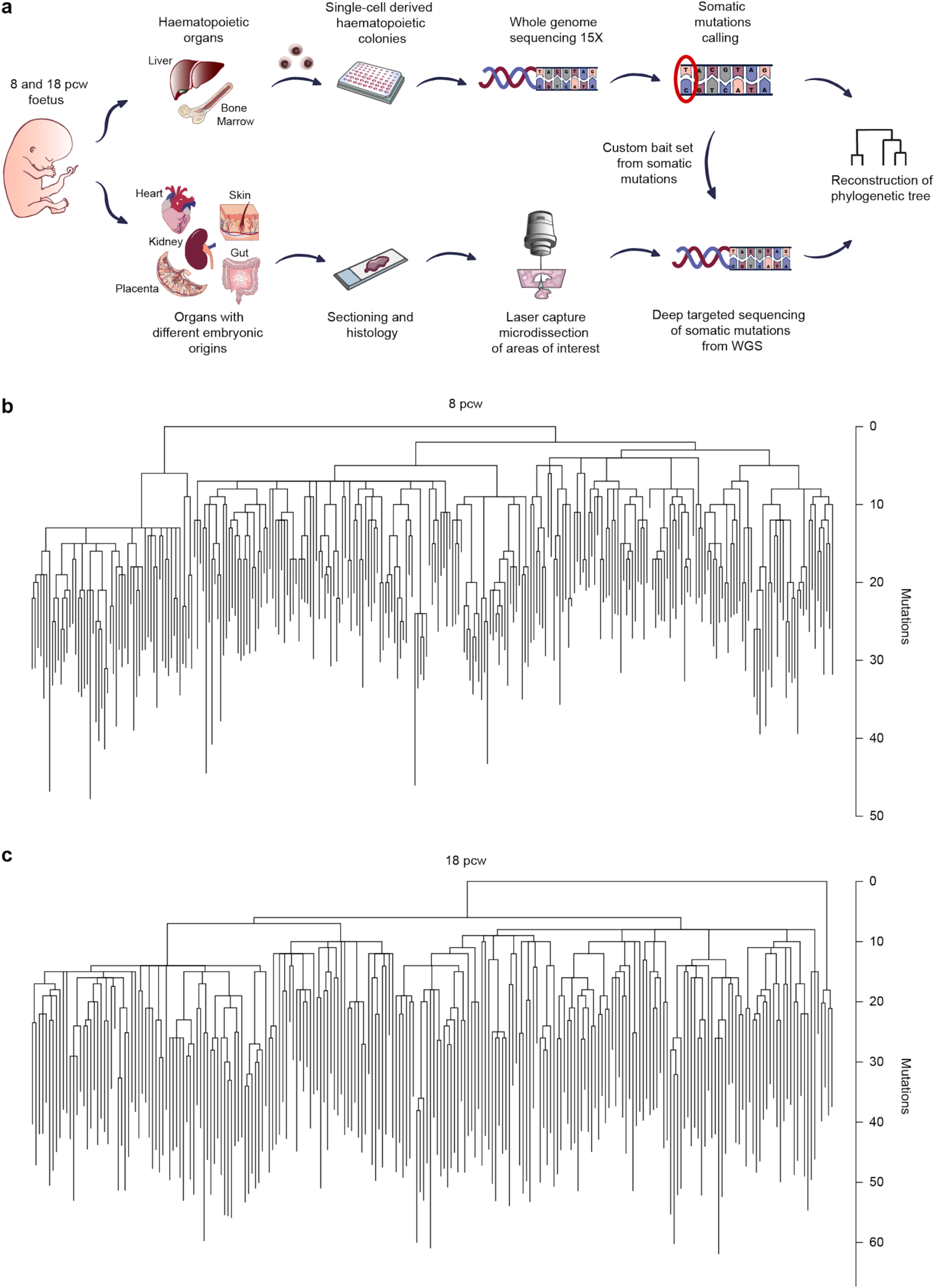
Experimental workflow and phylogenetic trees. **a**, Single HSPCs were isolated from an 8 pcw human foetal liver and from 18 pcw matched foetal liver and two femurs. Single-cell derived colonies were expanded in vitro and sequenced by WGS. Somatic mutations identified were used to reconstruct clonal relationships between cells. From the same two foetuses, non-haematopoietic organs with different embryonic origins were isolated, sectioned and laser-capture micro-dissected in order to produce well-defined sections with known developmental origin. These tissues underwent deep targeted sequencing using a DNA bait set covering the somatic mutations identified by WGS of blood colonies. Obtained data were integrated into the phylogenetic trees to highlight the time of separation of embryonic tissues during human development. **b**, Phylogeny of 277 single HSPCs from the liver of the 8 pcw foetus. **c**, Phylogeny of 234 single HSPCs from the liver and bone marrow of the 18 pcw foetus. Each tip of the tree is a single-HSPC derived colony. Cells sharing a common ancestor are grouped in clades deriving from a single node. Branch lengths are proportional to the number of acquired somatic mutations.

We first collected haematopoietic organs from two karyotypically normal male human foetuses of 8 and 18 pcw and used fluorescence-activated cell sorting (FACS) to isolate single HSPCs in liquid culture. Individual HSPCs were then expanded *in vitro* into colonies and sequenced by WGS to around 15× depth. From the liver of the 8 pcw foetus, we sequenced 277 colonies derived from single HSPCs (Lin-CD34+CD38+). At 18 pcw, both liver and bone marrow support blood production. Thus, we sequenced 234 colonies from phenotypically defined HSPCs (116 megakaryocyte-erythroid progenitors (MEPs), 105 common myeloid progenitors (CMPs) and 13 HSCs) from the liver (78 cells) and bone marrow of both femurs (79 and 77 cells) (Extended Data Fig. 1, Extended Data Table 1). The obtained sequencing depth (mean 22.6× for the 8 pcw colonies and mean 12.2× for the 18 pcw colonies) was sufficient to reliably call single-nucleotide variants (SNVs) (Extended Data Fig. 2). We applied stringent filtering criteria to remove germline mutations, recurrent artefacts from library preparation and sequencing, and mutations that were likely to have been acquired *in vitro*.

To validate the filtering strategy, we performed targeted resequencing of 501 of the 511 colonies to estimate the proportion of mutations that passed our filtering but were in fact subclonal or artefactual. Over 96% of mutations shared by more than one colony in the 8 pcw foetus and 100% of those in the 18 pcw foetus had variant allele fractions (VAFs) consistent with somatic mutations acquired *in vivo* (Extended Data Fig. 3a, b). Mutations passing filtering criteria were assigned to each cell. Mutations that were shared by multiple colonies were used to reconstruct phylogenetic trees of blood development (Fig. 1b, c). We constructed a tree in which the length of each branch is proportional to the number of somatic mutations accumulated over time within the inferred ancestral lineage. For clarity, here we refer to *lineage* as the direct descendants of a given ancestral cell.

In addition to HSPCs, we collected non-haematopoietic tissues of mesodermal, ectodermal, endodermal and extra-embryonic origin from the same two foetuses. We dissected the placenta, gut, kidney, heart and skin from the 8 pcw foetus; and kidney, heart and skin from the 18 pcw foetus. All tissues were fixed, paraffin-embedded, sectioned and cut using laser-capture microdissection (LCM) to obtain well-defined tissue structures (which we refer to as *microbiopsies* from now on) of specific extra-embryonic and embryonic origin. Microdissected tissues included the mesenchymal core from the placenta (deriving from the extra-embryonic mesoderm), syncytiotrophoblast (from the trophoblast), circulating blood (from the primitive wave of haematopoiesis), gut epithelium (from the endoderm), heart muscle (from the lateral plate mesoderm), kidney tubules (from the intermediate mesoderm), limb deep undifferentiated tissue (from mesoderm) intervertebral disc (from the paraxial mesoderm), epidermis (from the surface ectoderm) and skin peripheral nerve (from the neural ectoderm) (Extended Data Fig. 4). We then performed deep targeted sequencing (Extended Data Table 2) using custom DNA bait sets designed to include all shared and a majority of private somatically-acquired SNVs (please see Methods) detected in HSPC colonies by WGS.

### Acquisition of somatic mutations in early embryonic and foetal development

We estimated mutation burdens following adjustments for sensitivity and removal of *in vitro* mutations. In the 8 pcw foetus, the mean corrected SNV burden was 25 mutations per colony (standard deviation (sd) 6.1 mutations per colony) and in the 18 pcw foetus, the mean SNV burden was 42.6 mutations per colony (sd 7.7 mutations per colony) (Extended Data Fig. 5a). Interestingly, the observed standard deviations are broadly equivalent to that expected for a Poisson distribution, suggesting that all the lines-of-descent sequenced within each foetus have experienced the same mutation rates during development.

We detected a mean of 1.82 indels (range: 0 - 6) in the 8 pcw colonies and 1.23 (range: 0 - 5) in the 18 pcw colonies (Extended Data Fig. 5b). Due to the low numbers of indels, an equivalent sample-specific correction approach was not feasible. However, we used a Bayesian Poisson regression model to calculate a corrected mean indel burden of 2.09 indels per colony (95% confidence interval 1.90 - 2.29) for the 8 pcw foetus and of 2.13 indels per colony for the 18 pcw foetus (95% confidence interval 1.91 - 2.39). On average, the proportion of SNVs shared between multiple HSPC colonies was 0.62 for the 8 pcw foetus and 0.49 for the 18 pcw foetus (Extended Data Fig. 5c).

To further examine the genomic distribution of mutations identified by WGS we assessed the proportion of SNVs mapping to different genomic features. We identified a higher than expected proportion of SNVs in intergenic regions of the genome (Extended Data Fig. 6a). In contrast, we detected a lower than expected proportion of mutations in the introns, exons, and regions 5 kb upstream/downstream of the gene body (Extended Data Fig. 6b-c). This is consistent with previous reports of lower mutation rates in regions of transcription and accessible chromatin^14–16^.

The 96-profile mutation spectrum, which incorporates the trinucleotide context of the substitution, revealed a predominance of C>T transitions, primarily at CpG sites, which are associated with spontaneous deamination of methylated cytosine^17^. In both foetuses, the mutation spectrum of clonal mutations seen only in single colonies closely matched those of mutations shared by two or more colonies, confirming that the mutation spectrum does not noticeably change across the first half of gestation (Extended Data Fig. 7).

### Defining clonal relationships between foetal HSPCs

To determine clonal relationships between HSPCs, we constructed a phylogenetic tree for both 8 and 18 pcw foetuses (Fig. 1b, c). The root of a somatic lineage tree is, by definition, the zygote – it is likely that the initial bifurcation of the phylogeny therefore marks the first zygotic division to two blastomeres. Interestingly, the reconstructed phylogenetic trees revealed unequal contribution of each blastomere to the blood compartment (5:1 for 8 pcw and 59:1 for 18 pcw) indicating even higher asymmetry of embryonic contribution is possible for the first zygotic cell division than previously observed^13,18^.

The first 3 cell divisions (up to the 8-cell embryo) had a relatively high mutation rate, with a mean of 2.4 mutations acquired in each of the new daughter cells per cell division. This is comparable to estimates derived from VAFs of post-zygotic mutations in bulk sequencing^19^. From this point onwards in our two foetuses, there was a profusion of high-degree polytomies, defined as branch-points in the phylogenetic tree generating more than two descendants. In the context of a somatic lineage tree, a polytomy can only occur when there are cell divisions that generate no detected somatic mutations. Of the 5 cells from the 8-cell stage that contributed significantly to the HSPC population there are four polytomies (12-, 10-, 5-, 5-degree) and one dichotomy. Modelling of these polytomies suggests that the mutation rate decreases to less than 0.9 mutations/cell division at this stage of embryonic development (approximate Bayesian computation, see Methods). This coincides with zygotic genome activation in humans^20^, which may enable expression of DNA repair machinery.

### Timing the divergence of gut and blood during foetal development

In order to precisely determine the timing of individual mutation acquisition relative to key developmental events (i.e. trophectoderm formation, gastrulation and organogenesis), we deep-sequenced non-haematopoietic tissues for the somatic mutations that had been identified in HSPCs (mean tissue sequencing depth 450×, Extended Data Table 2). As we have seen, the mutation rate during embryogenesis is sufficiently informative that the phylogenetic tree essentially represents a high-resolution lineage tree defining the initial cell divisions of the embryo. A given mutation occurs in a specific embryonic cell, which can be placed accurately on the lineage tree – this mutation then acts as a permanent clonal mark inherited by all descendants of that cell. Using targeted sequencing of different tissues, with different developmental origins, we can identify mutations that are present in multiple tissue layers, organs or regions – such mutations must have occurred in cells that preceded the developmental split between those tissues. Therefore, the tissues that shared the fewest mutations with HSPCs were likely to be the first to separate from ancestral lines-of-descent that gave rise to the haematopoietic cells (*blood antecedents* from now on) and greater sharing of mutations in a given tissue suggests it diverged from blood antecedents later in development.

Using laser-capture microdissection, we generated multiple microbiopsies of non-haematopoietic tissues, such as the epithelium of the gut, which derives from the endoderm (Fig. 2a). DNA from the microbiopsies underwent targeted sequencing to characterise the presence or absence of each mutation, as well as the fraction of cells carrying each mutation in that microbiopsy.

**Fig. 2.**
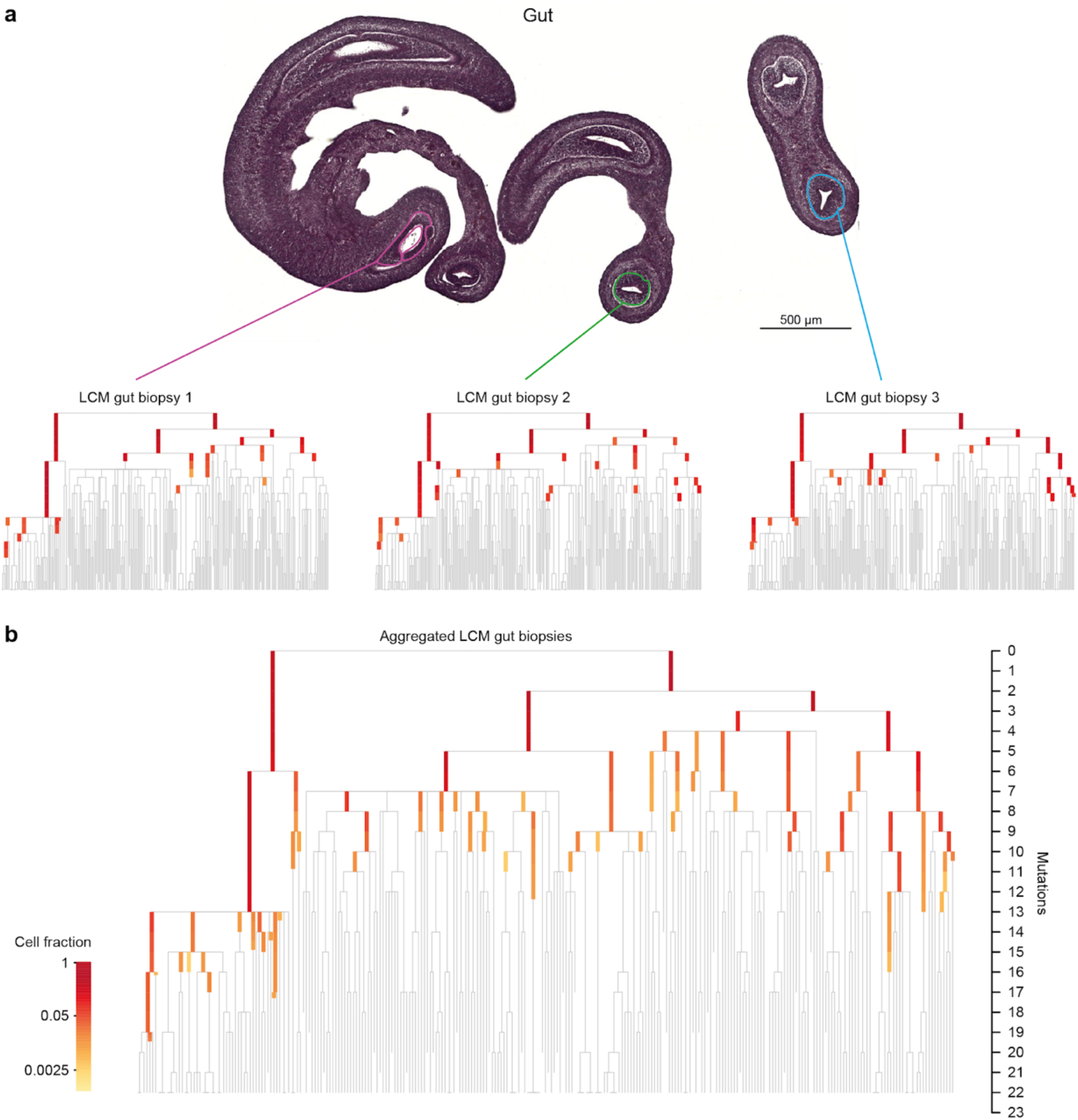
Reconstructing lineage divergence through targeted sequencing. **a**, Laser-capture microdissection was used to generate microbiopsies of 8 pcw gut epithelium. Somatic mutations shared with HSPCs were then detected by deep targeted sequencing. Mutations detected in HSPCs that were shared with gut epithelium are highlighted on the phylogenetic trees. The trees only show the first 22 mutations to highlight early branching. Three representative microbiopsy cuts are shown. Individual mutations in the phylogenetic tree are coloured on a grey-yellow-red log scale according to the fraction of cells in the biopsy carrying that mutation (with grey being absent), as shown in the scale in panel **b**. Mean sequencing depth: 89× for microbiopsy 1 (pink), 56× for microbiopsy 2 (green) and 91× for microbiopsy 3 (blue). **b**, Mutations shared after aggregating reads across 12 microbiopsies. Mean sequencing depth of aggregated microbiopsies: 841×. LCM, laser-capture microdissection.

For gut epithelium, we observed that mutations close to the root of the HSPC phylogenetic tree were universally present in the different microdissections, and contributed to high fractions of cells in the individual biopsies (Fig. 2a). From this, we can infer that the earliest cells in the embryo, such as those in the 4-cell and 8-cell cleavage stages, contributed large numbers of descendants to each of the gut microbiopsies (as well as blood HSPCs). As we traverse the phylogenetic tree from the root towards the tips, we found that the mutations became progressively less well represented in individual microbiopsies, before eventually becoming undetectable. Since each microbiopsy only samples a few hundred to thousand cells, and the average sequencing coverage was 70x per microdissection, we aggregated data from all 12 microbiopsies of gut epithelium to improve our statistical power for detecting mutations occurring later in embryogenesis (Fig. 2b).

The aggregated data for gut revealed that many lineages (≥60) made substantial contributions to both gut epithelium and blood HSPCs. These lineages must have predated gastrulation, when endoderm and mesoderm split from one another. Following individual lines-of-descent from root towards tip, different patterns emerged – for some, the cellular contribution of successive generations in that line-of-descent decreased steadily, suggesting that both daughter cells of each successive cell division contributed to gut epithelium. For others, the cellular contribution to gut epithelium stayed relatively constant over several generations, suggesting that only the daughter cells in that line-of-descent contributed meaningfully to gut epithelium.

### Timing specification of extra-embryonic and embryonic tissues

We proceeded in the same way, using LCM and deep targeted sequencing, to characterise all the embryonic and extra-embryonic tissues available to us (Extended Data Table 2). This enabled us to time the progressive divergence of different tissues from blood antecedents during embryonic development (Fig. 3a, b).

**Fig. 3.**
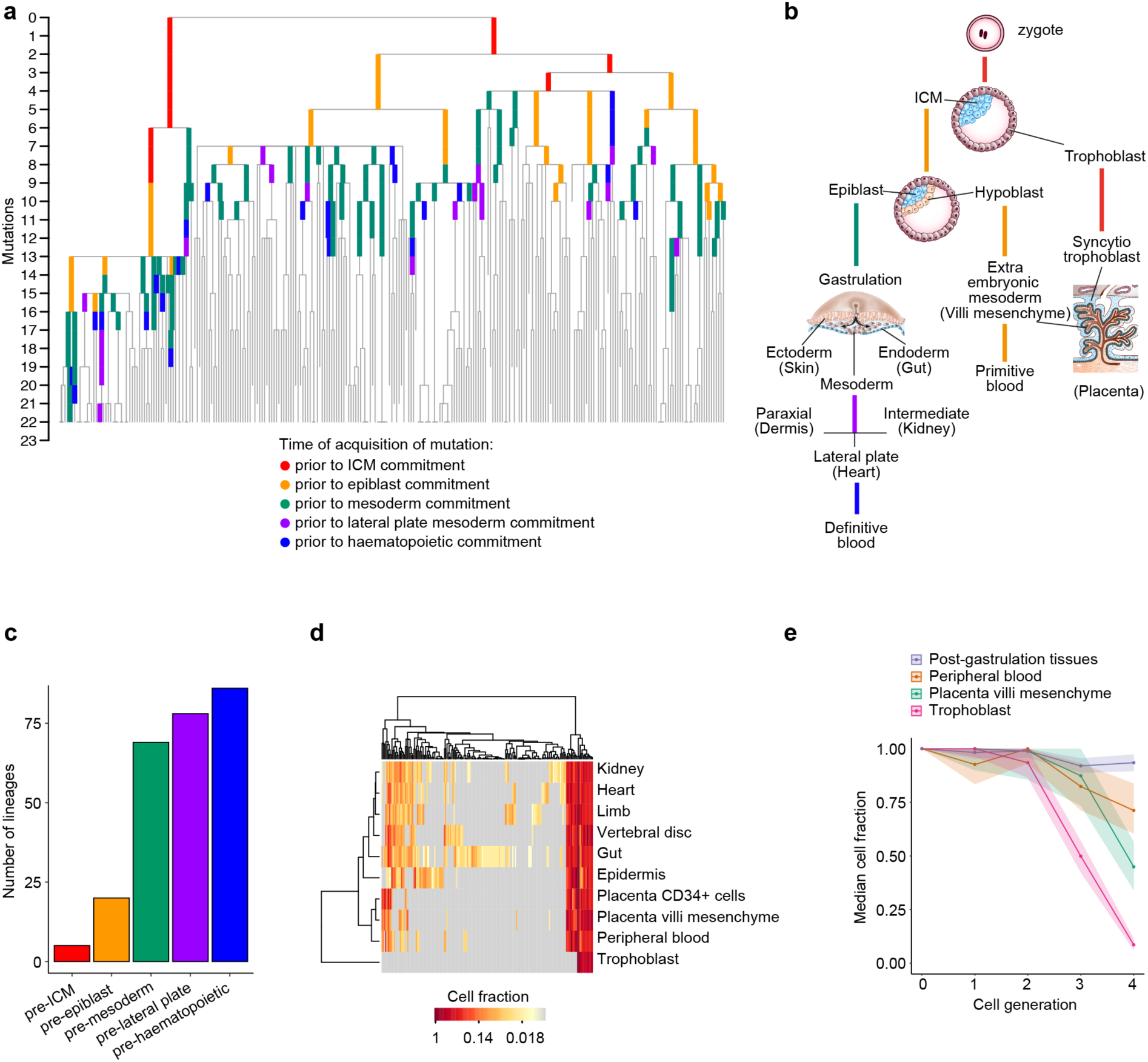
Timing of divergence of embryonic and extra-embryonic lineages during development. **a**, Phylogenetic tree showing the clonal relationships of 8 pcw HSPCs. Mutations identified in HSPCs by WGS that were also detected in non-haematopoietic tissues are coloured on the tree according to the earliest diverging tissue in which the mutation was reliably detected. Branch lengths are proportional to the number of mutations accumulated. The tree is cut at 22 mutations to highlight the early branching of the tree. **b**, Diagram of main human developmental events. Lineage separation is coloured as per the phylogenetic tree. In brackets are examples of tissues collected from each embryonic lineage. ICM, inner cell mass. **c**, Bar plot showing the minimum number of HSPC antecedents present at the time of divergence of different developmental lineages. **d**, Heatmap representation of the targeted sequencing data. Each column represents an individual mutation, and each row a single tissue. The colour shows the VAF of the mutation in that tissue with grey indicating that it was not detected. The tissues are clustered using soft cosine similarity (see methods). **e**, Line plot showing lineage loss of microdissected tissues at different times (represented as cell generations from the zygote). Shaded areas represent confidence intervals. ICM, inner cell mass.

In the mouse, the molecular distinction between trophoblast and ICM occurs at the 8-16 cell stage, but the exact timing in humans remains unknown^21^. In the 8 pcw foetus, we found that the trophoblast-derived tissue was the first to diverge from blood antecedents at the 4- to 16-cell stage embryo (corresponding to 2 to 4 generations from the zygote; *generation* is defined here as a round of cell division), thus following a timeline similar to mouse development. We can directly infer the presence of five distinct blood antecedents present at the time of trophoblast divergence in the embryo (Fig. 3c).

The next major event in embryogenesis is epiblast specification. We can infer that a minimum of 20 blood antecedent lineages already existed prior to epiblast specification (Fig. 3c). A recent study of *in vitro* human development up to the pre-gastrulation stage reported that specification of trophoblast, hypoblast and epiblast was evident by immunostaining of specific markers at 8 days post fertilisation. At this stage, the embryo comprised ∼500 cells, with ∼400 cells in the trophoblast, ∼60 in the epiblast and ∼40 in the hypoblast^22^. Therefore, there is a strong concordance between our estimates and the estimates based on immunostaining of epiblast-committed cells in embryos at the specific developmental stage. Our analysis, however, has the property that we can ascertain their ultimate contribution to organogenesis, not just their expression pattern at the time of specification. The concordance in numbers by the two methods does indeed suggest that the majority of cells expressing epiblast markers do in fact go on to contribute to detectable fractions of cells during organogenesis.

Proceeding further into embryonic development, we infer that at least 70 blood antecedents existed prior to mesoderm specification. Interestingly, the number of mutations shared between HSPCs and lateral, intermediate and paraxial mesoderm-derived tissues compared to endoderm- and ectoderm-derived tissues was similar, indicating that specification of each mesodermal component occurred within a few generations of mesoderm formation (Fig. 3a).

The origin of extra-embryonic mesoderm in humans has been debated for the last five decades, with mouse studies showing it derives from the primitive streak, while histological observations in humans point towards a hypoblast origin^6,77,8^. To address this issue, we compared the scope of shared mutations in the mesenchymal core of the placental villi and peripheral blood to that of tissues of known trophoblast and post-gastrulation origin. We observed that tissues derived from extra-embryonic mesoderm clustered separately from trophoblast-derived and post-gastrulation tissues (Fig. 3d and Extended Data Fig. 8a). In addition, 90% of trophoblast lineages were already lost in the tree by the fourth generation, in contrast to 25% and 50% of peripheral blood and placenta mesenchyme respectively and 10% of post-gastrulation lineages (Fig. 3e). Our data therefore support the hypothesis that the extra-embryonic mesoderm in humans emerges from the hypoblast^8^.

### Relative contributions of cells to germ layers at gastrulation

It is not known whether equal numbers of cells at gastrulation contribute to the tissues of all three germ layers. Interestingly, we found that the pattern of shared mutations in ectodermal tissues was distinct from that observed in mesodermal and endodermal tissues (Fig. 4). In non-haematopoietic mesodermal and endodermal tissues, we detected mutations from the majority of the blood antecedents up until 5-6 generations from the zygote, though each antecedent gave rise to a small proportion of cells from sequenced tissues at 8 pcw (Figure 4a, 4e and 4g). This implies that many epiblast cells contributed to mesoderm and endoderm development, and made broadly similar, but individually small, fractional contributions to these germ layers.

**Fig. 4.**
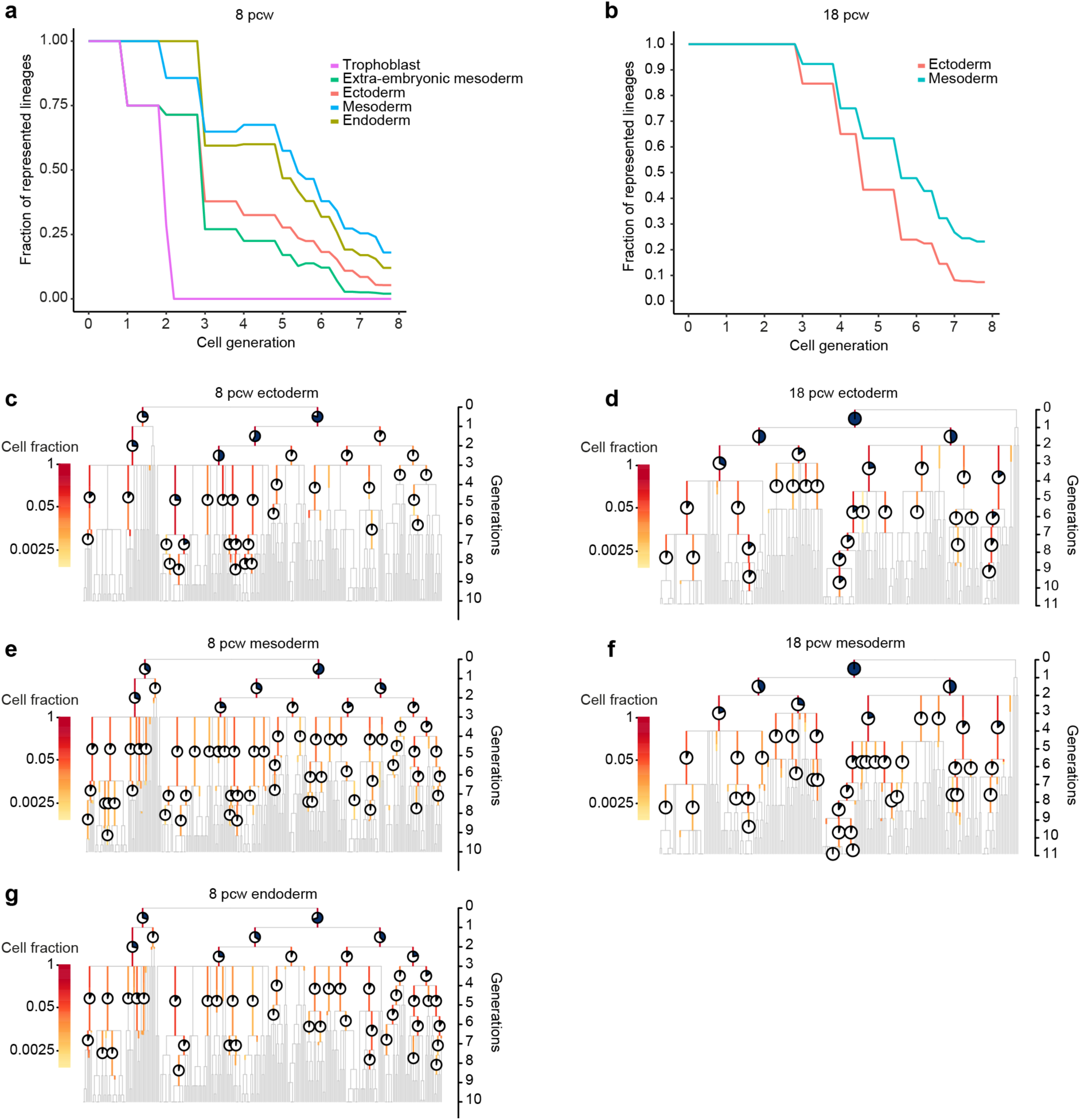
Representation of haematopoietic lineages in developmentally-defined non-haematopoietic tissues. **a**, Line plot showing the fraction of captured haematopoietic lineages in the 8 pcw phylogeny that are also detected in non-haematopoietic post-gastrulation tissues, grouped by germ layer. This is plotted over successive generations, during which the total number of lineages increases, but the proportion of lineages detected decreases. **b-d**, 8 pcw phylogenetic trees highlighting mutations detected in HSPCs that were shared with **b**, ectoderm-derived tissues, **c**, mesoderm-derived tissues and **d**, endoderm-derived tissues. The fraction of cells deriving from each lineage is plotted on a donut plot, with the black line representing the median cell fraction, and the dark and light blue segments representing the lower and upper 95% confidence intervals respectively. **e**, Line plot showing the fraction of captured haematopoietic lineages in the 18 pcw phylogeny that are also detected in non-haematopoietic tissues. For clarity the minor branch has not been considered as it does not contribute to any of the sequenced tissues. **f-g**, 18 pcw phylogenetic trees highlighting mutations detected in HSPCs that were shared with **f**, ectoderm-derived tissues and **g**, mesoderm-derived tissues. Trees in this figure were made ultrametric.

In ectoderm, however, mutations from fewer blood antecedents were detected, though with each giving rise to a relatively high cellular fraction of sequenced ectodermal microbiopsis (Figure 4b). Importantly, the individual lines-of-descent for ectoderm were detectable down to the same number of generations as for endoderm and non-haematopoietic mesoderm (typically 6-8 generations): there were just fewer of them. Moreover, following individual lines-of-descent from root to tip in the ectodermal deep sequencing showed that cellular contributions frequently stayed constant over several generations. This implies that the different pattern evident for ectoderm was not explicable by varying statistical power of mutation detection or the problem of lineages missing from our phylogeny. Instead, it suggests that relatively fewer cells from the epiblast contributed to the ectodermal tissues we sequenced than to endoderm or non-haematopoietic mesoderm.

A comparable pattern was seen in the 18 pcw foetus with two blood antecedents from around the ninth generation giving rise to a quarter of cells from sequenced ectodermal tissues. A similar limited set of blood antecedents predominated in all three ectodermal tissues that we sequenced (epidermis, hair follicle and peripheral nerve). While we cannot exclude the possibility that ectoderm antecedents migrate to a given anatomic region of the body, prior to their specification to distinct ectodermal derivatives, the more plausible conclusion would be that ectoderm originates from a smaller set of antecedents compared to endoderm and mesoderm.

### Trafficking of HSPCs across different haematopoietic sites

The colonisation of the long bones by haematopoietic cells from the liver starts around the 11th week of development^9^. From then on and up until birth, both the liver and the bone marrow are active sites of haematopoiesis. To investigate the kinetics of HSPC migration across early haematopoietic organs in 18 pcw foetus, we collected HSPCs from three separate sites of blood production. Out of the 234 HSPCs analysed, 78 were isolated from the liver, 77 from the bone marrow of femur 1 and 79 from femur 2. We did not observe statistically significant clustering (P=0.24) by organ of origin in the phylogeny, suggesting extensive traffic of haematopoietic stem and progenitor cells between liver and bone marrow (Extended Data Fig. 9a,b).

## Discussion

Our data reveal an initial high rate of somatic mutation acquisition, which drops markedly after 3 generations, coinciding with maternal-to-zygotic transition. Although at this stage of development, zygotic transcription is still inactive, the extensive chromatin accessibility^23,24^ and dilution of maternally-produced DNA-repair factors in dividing blastomeres could temporarily leave the genome more prone to accumulating mutations in the first generations. The reconstructed phylogenetic trees revealed an unequal contribution of each blastomere to the blood compartment which is in line with the stochastic nature of early cell fate decisions. This was particularly striking in the 18 pcw foetus, where just four out of 234 examined blood cells originated from one of the blastomeres. These four cells did not share any mutations with non-haematopoietic tissues of mesodermal and ectodermal origin (Extended Data Fig. 8) suggesting that the progeny of that blastomere had modest contribution to the embryonic tissues by either committing preferentially to trophoblast and hypoblast or dying out.

Based on the VAFs obtained from targeted sequencing in the 8 and 18 pcw foetus we further estimated that the separation between trophoblast and ICM occurs at the 8- to 16-cell stage with five distinct blood antecedents being present at the time. The number of blood antecedents in the embryo increased to 20 at the epiblast stage and a minimum of 70 at the time of mesoderm specification. Interestingly, our data suggest that ectoderm originates from a smaller set of blood antecedents compared to endoderm and mesoderm. Once the mesoderm was formed the specification of mesodermal components (i.e. lateral, intermediate and paraxial mesoderm) occurred within a few generations. Finally, we show that the primitive blood, captured in the circulation of the 8 pcw foetus, originates from hypoblast, as demonstrated by its higher level of mutation sharing with the extra-embryonic mesodermal tissues compared to tissues of known post-gastrulation origin. In this study we provide the conceptual framework for the future exploration of the outstanding questions in human development through the use of naturally occurring somatic mutations as barcodes.

## Acknowledgements

The authors would like to thank the Wellcome Sanger Institute (WSI) Cytometry Core Facility for their help with single-cell index sorting, the WSI DNA pipelines for their contribution in sequencing the data and the WSI Cancer, Ageing and Somatic Mutation programme IT and sample teams for their bioinformatic and logistical support. We would also like to thank Jana Eliasova for her precious support and help with the illustrations and the Human Developmental Biology Resource (HDBR) for providing samples.

## Authors contributions

A.C., A.M.R. and P.C. conceived the study; A.M.R. performed all the experiments with help from B.M; M.S.C. carried out the computational analysis of the WGS data, under the supervision of P.C.; Y.H. prepared all histology sections for microdissection; T.B. assisted with the targeted sequencing strategy; T.B and P.R. assisted with the laser capture micro-dissections; E.H. and L.M. assisted with annotating histology slides; N.W., T.C. and E.M. assisted with somatic mutation calling from WGS data, the construction of the phylogeny and the assignment of mutations to the tree; N.W., J.N. and K.D. helped design and implement population simulation models to help interpret the phylogenies; A.C., A.M.R, and M.S.C. designed the figures and wrote the manuscript with inputs from the other authors. All authors approved the final version of the manuscript.

## Methods

### Ethics and tissue acquisition

Tissues used in this study were isolated from two karyotypically normal male human foetuses (8 and 18 pcw) following termination of pregnancy and informed written consent. All material was provided by the Joint MRC/Wellcome Trust (Grant MR/R006237/1) Human Developmental Biology Resource (http://www.hdbr.org), in accordance with ethical approval by the NHS Research Health Authority, REC Ref: 18/LO/0822. All tissues were processed on the day of collection.

### Generation of single-HSPC colonies

Foetal liver tissue was passed through a 70 **μ**m strain with cold PBS (Gibco) to generate a single-cell suspension. Bone marrow was isolated by flushing cold PBS into the cavity of the two femurs (for the 18 pcw sample) and passed through a 70 **μ**m strain to generate a single cell suspension. Cells were centrifuged at 300 g for 5 minutes at 4°C and resuspended in Red Blood Cell lysis buffer (eBioscience) for 2 minutes. The lysis reaction was stopped by adding 20 ml of cold PBS. MACS columns (Miltenyi Biotec - 130-090-101) were used for live cell enrichment, following the manufacturer’s instructions. For the 18 pcw tissues, a CD34+ cell enrichment step was also performed, using MACS columns (Miltenyi Biotec - 130-046-702) following the manufacturer’s instructions. Cells were stained with antibodies in a total volume of 100 **μ**m PBS with 5% FBS (Gibco) for 30 minutes covered from light at 4°C and filtered into polypropylene FACS tubes (ThermoFisher) to a final volume of 500 **μ**l of PBS with 5% FBS. Single HSPCs were index-sorted with a BD Influx Sorter in liquid culture (as previously described^25^) into 96-well plates seeded with MS5 feeder layer (DSMZ) and cultured for 6-8 days at 37° and 5% CO_2_. At the end of the culture, cells were collected and filtered into polypropylene FACS tubes. Cells were sorted with a BD Influx Sorter into 96-well plates and lysed using the Arcturus PicoPure Kit (Applied Biosystems) according to the manufacturer’s instructions.

### Tissue processing for LCM

Tissues from the 8 pcw (placenta, heart, kidney, gut, skin and vertebrae) and 18 pcw (skin, kidney and heart) samples were fixed in PAXgene Tissue FIX (Qiagen) for 4 and 24 hours respectively and embedded in paraffin using a Tissue-Tek VIP machine (Sakura). Sections of 5-10 **μ**m were cut using a microtome (Leica-RM2265), placed on PEN MembranSlide glass slides with a 2.0**μ**m membrane (Leica) and dried in a laboratory oven (Genlab) at 37°C. Sections were dehydrated with a series of ethanol (VWR) washes and stained with Mayer’s haematoxylin (Sigma) / Accustain Eosin Y (Sigma) before a final xylene (Sigma) cleaning step. Slides were dried overnight. High resolution imaging was performed using a slide scanner (Hamamatsu). Areas of interest were annotated on tissue scans and isolated with a microscope equipped with a laser-capture microdissection system (Leica LMD7). Microbiopsies were collected in 96-well plates and lysed using the Arcturus PicoPure Kit (Applied Biosystems) according to the manufacturer’s instructions.

### Library preparation and sequencing

Library preparation for both the HSPC colonies and LCM microbiopsies was performed as previously described^26^. Paired end sequencing reads (150bp) were generated using either the Illumina NovaSeq® 6000 platform (8 pcw foetus) or Illumina HiSeq® 4000 (18 pcw foetus) resulting in ∼22.6× coverage and ∼12.2× coverage per colony respectively. BWA-MEM was used to align sequences to the human reference genome (NCBI build37).

### Mutation discovery and tree-building from whole-genome sequencing data

Whole-genome sequencing data from single-cell colonies demonstrated variable levels of contamination by mouse DNA from the mouse feeder cell layer. Therefore, sequencing reads were filtered using the ‘Xenome’ algorithm which tests each read for mapping against the human and mouse reference genomes, and removes reads of clear mouse-origin, or that are ambiguous^27^. Remaining reads were aligned to the human reference genome and single-nucleotide variants (SNVs) and indels were called against an unmatched reference genome using the in-house pipelines CaVEMan and Pindel^28,29^. For all mutations passing quality filters in at least one sample, in-house software (cgpVAF, https://github.com/cancerit/vafCorrect) was used to produce matrices of variant and normal reads at each site for all HSPC colonies from that foetus.

Multiple post-hoc filtering steps were then applied to remove germline mutations, recurrent library prep and sequencing artefacts, and probable in vitro mutations, as detailed below:

1. A custom filter designed to remove artefacts specifically associated with the library prep process used for low input DNA concentrations (https://github.com/MathijsSanders/SangerLCMFiltering). Specifically, this includes those introduced by cruciform DNA structures.
2. A binomial filter was applied to aggregated counts of normal and variant reads across all samples. Sites with aggregated count distributions consistent with germline single nucleotide polymorphisms were filtered.
3. A beta-binomial filter was applied to retain only mutations whose count distributions across samples came from an over-dispersed beta-binomial distribution consistent with an acquired somatic mutation.
4. Mutations called at sites with abnormally high or low mean coverage were considered unreliable/ possible mapping artefacts and were filtered.
5. For each mutation call, normal and variant reads were aggregated from positive samples (≥ 2 variant reads). Sites with counts inconsistent with a true somatic mutation were filtered.
6. Remaining mutations were only retained if there was at least one sample that met all minimum thresholds for variant read count (≥3 for autosomes, ≥2 for XY chromosomes), total depth (≥6 for autosomes, ≥4 for XY chromosomes), and a binomial probability of >0.1 that variant:normal counts reflect that expected for a genuine somatic mutation (0.5 for autosomes and 0.95 for XY chromosomes [not set to 1 to allow for artefacts]).

Filter threshold levels for filter steps 1-4 were set by using a combination of past experience, review of threshold parameter histograms, and experimentation. Mutational signatures and consistency with the phylogenetic tree were used for validation. In general, more stringent thresholds were used than in analyses on adult samples, due to the low number of anticipated true positives.

### Phylogenetic tree construction

For purposes of tree building, samples were assigned a genotype for each remaining mutation. All samples with ≥2 variant reads and a probability > 0.05 that counts came from a distribution expected for a somatic mutation were genotyped as present. Samples with 0 variant reads and a depth of ≥6 at a given site were genotyped as absent. Samples not meeting either of these criteria were genotyped as ‘unknown’.

A genotype matrix of shared mutations (a positive genotype ≥1 sample [i.e. that would be informative for tree-building]) was fed into the MPBoot program^30^. This can incorporate ‘unknown’ genotypes, and constructs a maximum parsimony phylogenetic tree with bootstrap approximation.

### Mutation assignment back to the phylogenetic tree

All mutations retained post-filtering were then assigned back to individual branches on the phylogenetic tree using the in-house developed R package ‘treemut’ (https://github.com/NickWilliamsSanger/treemut). This goes back to the original count data, and uses a maximum-likelihood approach to assign each mutation to a specific branch. Edge lengths are then made proportional to the number of mutations on the branch.

### Inference of the mutation rate after the 8-cell-stage embryo

After the third round of cell division (corresponding to the 8-cell-stage embryo), the 8 pcw HSPC phylogeny shows a profusion of high degree polytomies. Specifically, of the five blood antecedents from the 8-cell stage for which we have significant numbers of descendants, there are 12-, 10-, two 5-degree polytomies and a single dichotomy. The true degree of these polytomies may be underestimated if descendant lineages are not captured in the HSPC phylogeny. We ran 1000 simulations of 5 independent cells dividing and acquiring mutations randomly according to a Poisson distribution with lambda corresponding to a specific average mutation rate, and recorded the proportion of simulations that resulted in polytomies at their roots of *at least* the degrees observed. We repeated these simulations for average mutation rates ranging from 0.5 to 1.5 mutations per cell division. These simulations showed that once the mutation rate rose to 0.9 mutations per cell division, ≤5% of simulations gave results consistent with the data, and once the mutation rate was 1.0, this proportion dropped to ≤1%.

### Targeted sequencing

Individual custom DNA capture panels were designed for each foetus according to the manufacturer’s guidelines (SureSelect^XT^ Custom DNA Target Enrichment Probes, Agilent). For the 8 pcw foetus a 0.85 Mb panel was designed to capture regions of all SNVs and indels passing the previously described filtering steps. For the 18 pcw foetus a 0.5 Mb bait set was designed to capture SNVs only, and included all shared SNVs and around half of private SNVs. No repeat-masking was used in the design process, and hence on-target read percentage was low (<10% for both panels).

### Validation of somatic mutations called in haematopoietic colonies

DNA libraries from the WGS-sequenced 8 pcw and 18 pcw HSPC colonies were combined into pools of 24-32 samples, hybridized to the appropriate capture panel, multiplexed on flow cells, and subjected to paired-end sequencing (150-bp reads) on the HiSeq4000 machine (Illumina). BWA-MEM was used to align sequences to the human reference genome (NCBI build37). cgpVAF was used to compile variant read counts and total read depth from all included mutant loci. For each mutation the total variant read and total depth counts were aggregated for all colonies expected to carry the mutation, based on the phylogeny structure.

For the 18 pcw foetus, 2137 private SNVs were included in the bait set. Of these, 1770 achieved a depth of ≥8× in the colony in which they were originally called (the “minimum depth” set); 273 achieved a depth of ≥40× in the colony in which they were originally called (the “high depth” set). The read counts from each set were used as input to a binomial mixture model to ascertain the proportion of mutations that were clonal (and therefore likely to be true somatic mutations), sub-clonal (and therefore *in vitro* mutations), or absent. The binomial mixture model uses an expectation maximisation algorithm to calculate the best combination of between 1 and 4 binomial distributions to fit the observed data. For each individual mutation it assigned the most likely binomial distribution for it to be derived from i.e. was it most likely clonal or sub-clonal. Similar results were obtained with both the “minimum depth” and “high depth” mutation sets (Extended Data Fig. 3).

For the 8 pcw foetus, 3274 private SNVs were included in the bait set. Of these, 1431 achieved a depth of ≥8× (minimum depth set) in the colony in which they were originally called, and 34 a depth ≥40× (high depth set). Otherwise analysis proceeded similarly as per the 18 pcw HSPC colonies.

Indels were included in the 8 pcw capture panel only. There were 246 indels in total: 196 private and 50 shared. Of these, 73 private and 33 shared indels on autosomes achieved a total depth of ≥8× in colonies in which they were called. Applying the binomial mixture model to these read counts showed that counts for both private and shared mutations were consistent with a single binomial distribution with probabilities of 0.49 and 0.475 respectively.

### Detection of somatic mutations discovered in HSPC colonies in non-haematopoietic tissues

DNA libraries from LCM cuts of non-haematopoietic tissues were combined into pools of 16 samples, hybridized to the appropriate capture panel, multiplexed on flow cells, and subjected to paired-end sequencing (150-bp reads) on both the HiSeq4000 and NovaSeq 6000 machines (Illumina). For the 68 DNA libraries from the 8 pcw foetus LCM cuts, one HiSeq4000 lane and two NovaSeq 6000 lanes were used. For the 48 DNA libraries from the 18 pcw foetus, two HiSeq4000 lanes were used. BWA-MEM was used to align sequences to the human reference genome (NCBI build37). cgpVAF was used to compile variant read counts and total read depth from all included mutant loci.

Mutation calling was performed using a depth-aware naive Bayesian classifier comparing the two competing models:

H_0_: Observed VAF is consistent with sequencing error; versus

H_A_: Observed VAF is consistent with the mutation being truly present in the sample.

The data comprised counts of total depth, *n*_*i,j*_, and number of reads, *x*_*i,j*_, for the *i*th mutation in the *j*th sample. We estimated the sequencing error rate for the *i*th mutation as coming from a *BetaBinomial*(*α*_*E*_, *β*_*E*_ *n*_*i,j*_) distribution, using deep sequencing data from colonies that were known to be negative for the mutation by their position on the phylogenetic tree. Estimates were derived using the method of moments, empirically weighted for differences in sequence depth across colonies as described by Kleinman^31^. The data for a true mutation were assumed to be drawn from a *BetaBinomial*(*α*_*M*_, *β*_*M*_, *n*_*i,j*_) distribution with *α*_*M*_ = 1.5; *β*_*M*_ = 10. The prior probability of a mutation being present in a tissue, *π*_*i,j*_, was set as the proportion of HSPC colonies from the WGS phylogeny harbouring that mutation. Using this and the usual beta-binomial probability density function, then, the posterior probability for H_A_ is: 

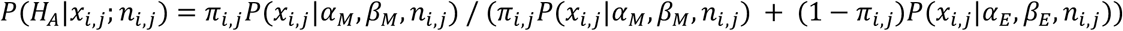

For aggregated tissues, the posterior probability of the mutation being present in the tissue as a whole was calculated as one minus the product of the probabilities of the mutation being absent in each individual microbiopsy. The knowledge of the phylogeny structure was then used to clean up calls that did not concord with mutation calls from neighbouring branches. Mutations that were called as present from distal branches when their ancestral branches were absent were removed; mutations which had been called as absent, but had confident mutation calls from both distal and proximal branches (or from the same branch) had their posterior probabilities elevated. This latter scenario usually occurred for mutations where the depth was unusually low and no positive reads had been found in a given tissue. Trees for figures 3 and 4 were then plotted with the VAF of individual mutations called as present overlayed on the underlying HSPC phylogeny.

### Inference of numbers of lineages at different developmental stages

The presence of mutations from the HSPC phylogeny in non-haematopoietic tissues demonstrates that the mutation was acquired in a blood antecedent cell that still had the capacity to differentiate into both blood and the non-haematopoietic tissue in question. By considering a phylogeny including only those branches shared with a specific non-haematopoietic tissue you can visualize the phylogeny of these multipotent cells. The number of descendant non-shared branches stemming directly from this shared phylogeny is a direct read of the minimum number of more committed HSPC antecedents arising from these multipotent antecedents.

We performed this analysis considering the shared phylogenies of non-haematopoietic tissues that developmentally diverge from blood at different points:

1. the trophoblast-shared phylogeny: the inner cell mass (ICM) /trophoblast divergence,
2. the mesenchymal core-shared phylogeny: the epiblast/ hypoblast divergence,
3. the endoderm- or ectoderm-shared phylogeny: the mesoderm /non-mesoderm divergence,
4. the kidney- or limb-shared phylogeny: the lateral-plate/ non-lateral-plate mesoderm divergence,
5. the heart-shared phylogeny: the haematopoietic lateral-plate mesoderm/ non-haematopoietic lateral-plate mesoderm divergence.

For the early lineage commitment events (ICM commitment and epiblast commitment) our HSPC phylogeny captures the vast majority of antecedents from these time points, and therefore the estimates are likely close approximations of the true values. For the later commitment events, the phylogeny only captures a small fraction of antecedents, and therefore the numbers represent lower bounds on the true value.

### Clustering of targeted sequencing results from non-haematopoietic tissues

To evaluate the similarity of the pattern of shared mutations and their VAFs in the different non-haematopoietic tissues we used the soft cosine similarity statistic, a measure used extensively in natural language processing. In this measure, the similarity between two sets of observations is defined as: 

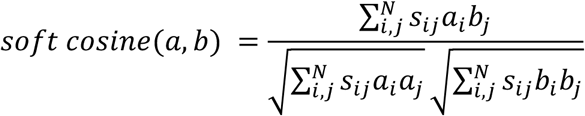

Here the *i* and *j* represent an iteration over all mutations; the *a*_*i*_ and *b*_*j*_ represent the natural log of the VAF of mutation *i* in tissue *a* versus mutation *j* in tissue *b* respectively; the *s*_*ij*_ represents a similarity score for mutation *i* and *j* which for our analysis we considered to be the inverse of the 1 + the minimum total branch distance between the two mutations.

The euclidean distance between tissues was then calculated, and the tissues clustered by hierarchical clustering using the *hclust* function from the “stats” package within R. The calculated dendrogram is displayed alongside the heatmap in Figure 3d.

### Measure of lineages lost up to the 4th generation from the zygote

In the 8pcw foetus, we attempted to assign individual mutations to the round of cell division during which they originated. For the first two generations from the zygote (up to the 4-cell stage embryo) it was straight-forward to directly infer this with some confidence because (1) divisions are invariably associated with mutations due to the relatively high mutation rate i.e. they are dichotomies, and (2) the phylogeny captured many descendants from each of the 4-cell stage antecedents allowing each branch to be resolved. The fact that the VAFs across these branches sum to 0.5 across both embryonic and extra-embryonic tissues adds confidence that no lineages have been missed. For the 3rd generation (i.e. the 8-cell stage embryo) this was no longer true, as some of the 8-cell-stage antecedents contribute only to trophoblast. However, targeted sequencing data from matched trophoblast allowed an apparent single branch to be resolved into two separate branches resulting from 2nd and 3rd generation divisions. From the 4th generation, there are two major difficulties in assigning mutations to generations: (1) due to the drop in mutation rate, many divisions occur without the acquisition of new mutations making it impossible to know which of the polytomous branches is a result of which generation (2) we have fewer descendants from each lineage, meaning that each daughter branch may have mutations resulting from many successive cell divisions. For the purposes of figure 3e, we considered all daughter branches of 3rd generation antecedents as potential 4th generation divisions, although in reality they are a mixture of 4th and higher divisions. To resolve branches that had mutations from multiple successive cell divisions we again used the targeted sequencing data: a binomial mixture model was applied to aggregated read counts from all non-haematopoietic tissues, analysing mutations from each branch in turn. Branches with read counts best explained by more than one binomial distribution (i.e. containing mutations present in different cell fractions) were annotated such that only mutations from the highest probability binomial distribution were considered as 4th generation mutations.

Once mutations were assigned to each generation, the total cell fraction captured by our HSPC phylogeny for mutations acquired in the 1st - 4th cell divisions was calculated by summing the cell fractions for each branch in a given generation. The uncertainty in these cell fractions was calculated by bootstrapping the read counts from each branch 1000 times, repeating the sum of cell fractions across branches for each bootstrap, and obtaining the 0.025 and 0.975 quantiles from the resulting distribution for a 95% percentile interval.

### Construction of the phylogenies with branch lengths scaled by generations

As described above, assignment of branches to specific generations is relatively straight-forward for the first three generations. However, this becomes more challenging once there are frequent polytomies and fewer descendant cells from each branch.

To create an approximation of a “generation” tree where branches are placed along the y axis according to the generation in which the mutations arose we did the following:

1. Branches were manually assigned to generations as previously for the first 3 generations
2. Branches from polytomies were extended according to log_2_ of the number of descendants e.g. for an 8-degree polytomy, each branch would be extended to length 3, the average number of generations. In reality, in this example, individual branches may result from 1 to 7 generations from the antecedent, depending on the true phylogeny structure underlying the polytomy.
3. Terminal branches were extended to make the tree ultrametric.

For the analysis of the lineages represented through different generations (Figure 4a-b), single branches from polytomies were therefore considered as belonging to multiple generations. This approach means that the absolute proportions of represented lineages by generation are inaccurate as different assumptions will falsely increase/ decrease these proportions. However, for comparison between tissues it remains a useful measure.

### Analysis of mutation distribution in the genome

To evaluate whether the mutation distribution across different genomic features was comparable to that expected by chance we compared our data to a simulation of random mutagenesis within the human genome.

In this simulation the *bedtools random* command was used to generate 160,000 random sites within the human genome. From these, 5000 were subsampled such that the 32 possible trinucleotide contexts were represented in the same proportions as in the data. The central pyrimidine within each trinucleotide was then virtually “mutated” to an alternative base, again according to probabilities matching the mutational signatures of the data to generate a vcf file of 5000 mutational-signature matched mutations. The resulting genomic locations were annotated using the VAGrENT algorithm (https://github.com/cancerit/VAGrENT) whereby each mutation is assigned a Sequence Ontology term to classify its consequence.

This simulation was repeated 500 times and the distribution of the resulting proportions of mutations in different Sequence Ontology categories was compared to those observed in the true mutation sets.

### Analysis of molecular variance to look for phylogenetic clustering by anatomic location

Analysis of molecular variance (AMOVA) was used to test for phylogenetic clustering by anatomical location in the 18 pcw foetus. We compared HSPCs that had been isolated from the liver, femur 1, and femur 2.

The phylogeny was first made ultrametric using a bespoke method that gives more weight to the lengths of early branches. The mutational distance between any two HSPC colonies was then calculated (this is the minimum mutational “walk” to get from one sample to the other) for all possible colony-colony pairs. The sum of squared distances for “within location” and “between location” pairs was calculated, divided by their degrees of freedom, and this was used to obtain an estimate of the proportion of variation explained by differences in cell locations (the F statistic).

P values were obtained by randomly re-labelling the cell locations 30,000 times and calculating the same statistic. The given p value is therefore a proportion of the random re-assignments that had a more extreme F statistic than that observed in the data.

## Code availability

All scripts and some derived data sets are available at https://github.com/mspencerchapman/Phylogeny_of_foetal_haematopoiesis.

## Data availability

Whole genomes and targeted sequencing data will be available via European Genome-Phenotype Archive (EGA, https://www.ebi.ac.uk/ega/). Accession numbers not yet assigned.

## Extended Data

**Extended Data Fig. 1.**
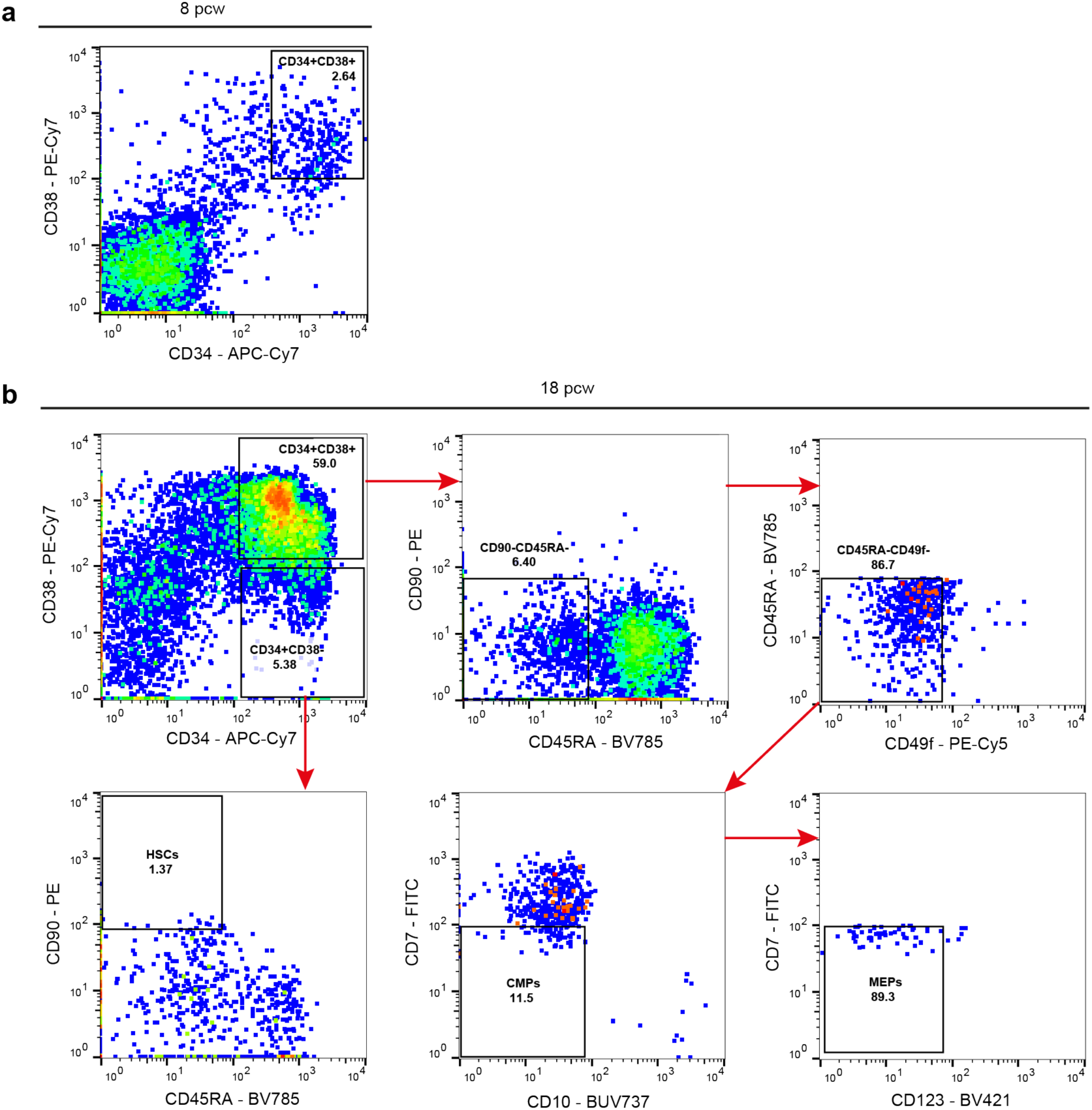
FACS sorting strategy for haematopoietic progenitor cells from foetal liver and bone marrow. **A**, Following the exclusion of debris and cell doublets by gating, anti-CD34 and anti-CD38 staining was used to sort single haematopoietic progenitor cells from the liver of a 8 pcw foetus. **b**, Sorting strategy for different stem and haematopoietic progenitor populations from matched liver and two femurs of a 18 pcw foetus. Following exclusion of debris and cell doublets, anti-CD34, anti-CD38, anti-CD90, anti-CD45RA, anti-CD49f, anti-CD7, anti-CD10 and anti-CD123 staining was used to sort haematopoietic progenitor populations. HSCs = haematopoietic stem cells, CMPs = common myeloid progenitors and MEPs = megakaryocyte-erythroid progenitors. n = 20,000 cells.

**Extended Data Fig. 2.**
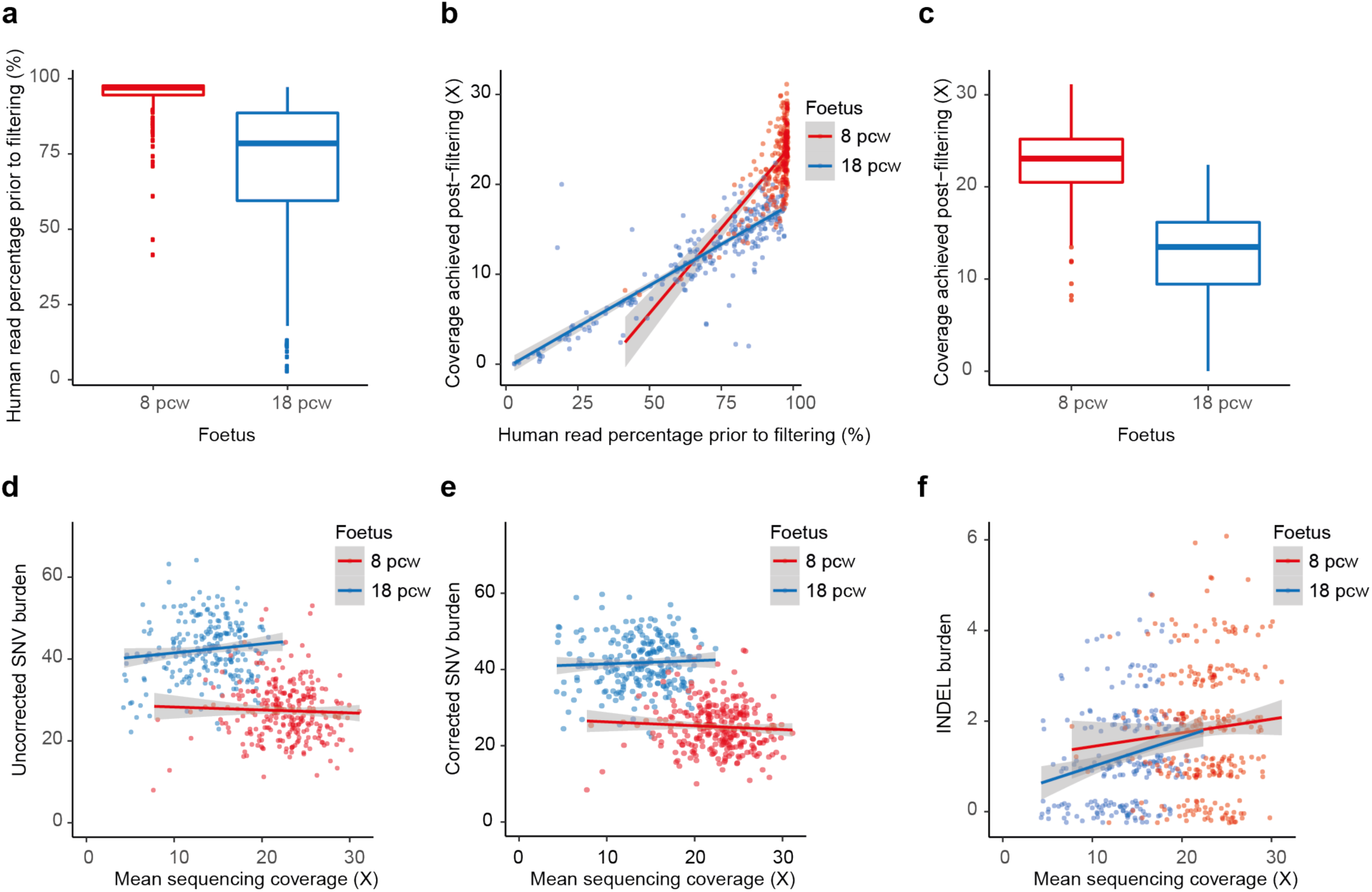
Sample contamination and sequencing coverage. **a**, Box plot showing the percentage of human sequencing reads, prior to the exclusion of contaminating mouse reads from the feeder layer. **b**, Dot plot showing, for each colony of the two foetuses, the final sequencing coverage, post-exclusion of mouse reads, against the percentage of human reads. **c**, Box plot showing final sample coverage for the two foetuses. **d**, Dot plot showing the uncorrected SNV burden per colony against sample coverage. **e**, Dot plot showing the corrected SNV burden per colony against sequencing coverage. **f**, Dot plot showing the uncorrected indel burden per colony against sequencing coverage. SNVs = single nucleotide variations.

**Extended Data Fig. 3.**
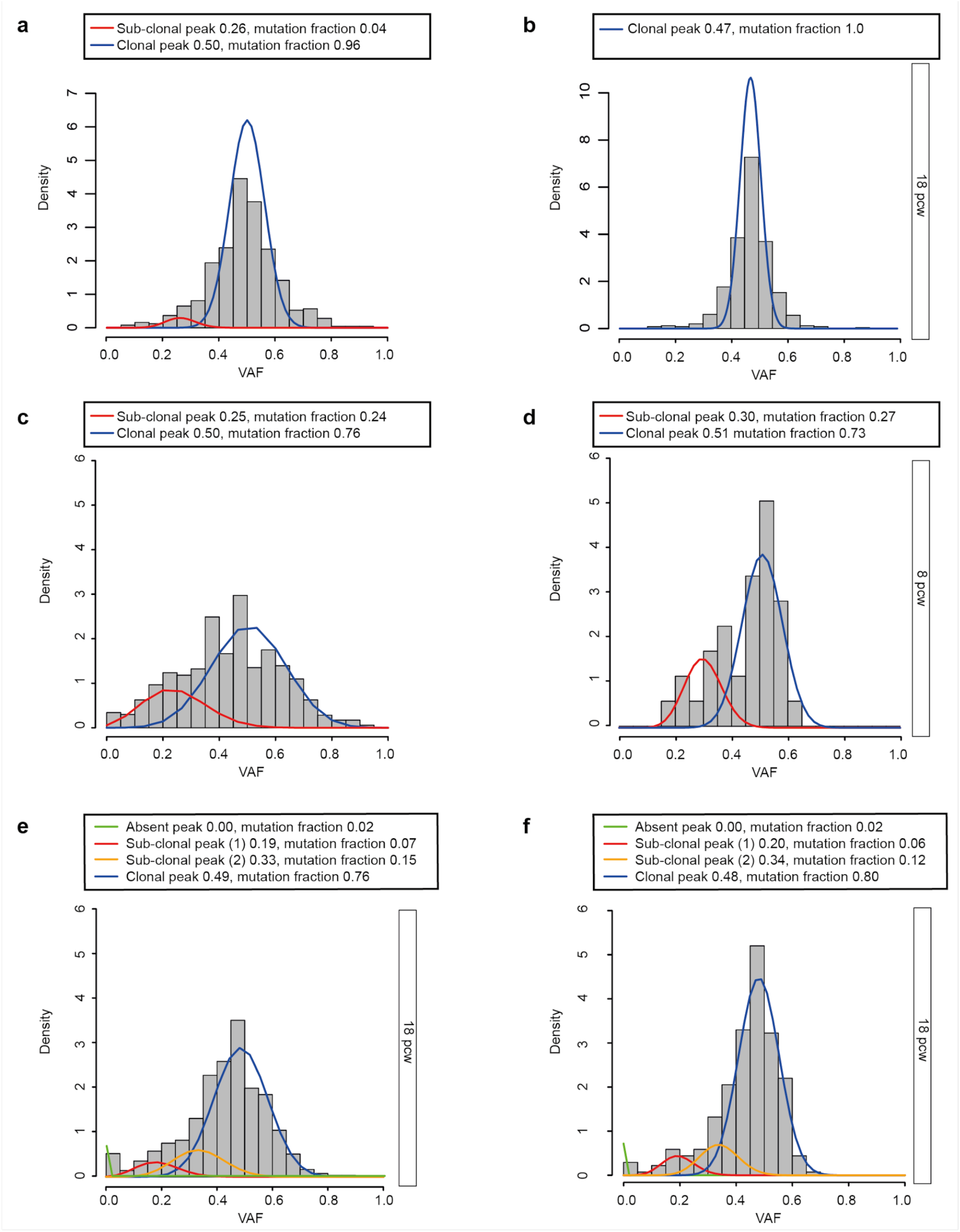
Mutation validation. Histograms showing variant allele fraction (VAF) of **a**, shared mutations with targeted sequencing depth ≥8× in the 8 pcw foetus, **b**, shared mutations with targeted sequencing depth ≥8× in the 18 pcw foetus, **c**, private mutations with targeted sequencing depth ≥8× in the 8 pcw foetus, **d**, private mutations with a high read depth ≥40× in the 8 pcw foetus, **e**, private mutations with depth ≥8× in the 18 pcw foetus, **f**, private mutations with a high read depth ≥40× in the 18 pcw foetus. Almost all shared mutations fitted a single clonal distribution. Private mutation distributions were best explained by a mixture of clonal (blue line, probability ∼0.50) and one or more sub-clonal binomial distributions (red and yellow lines).

**Extended Data Fig. 4.**
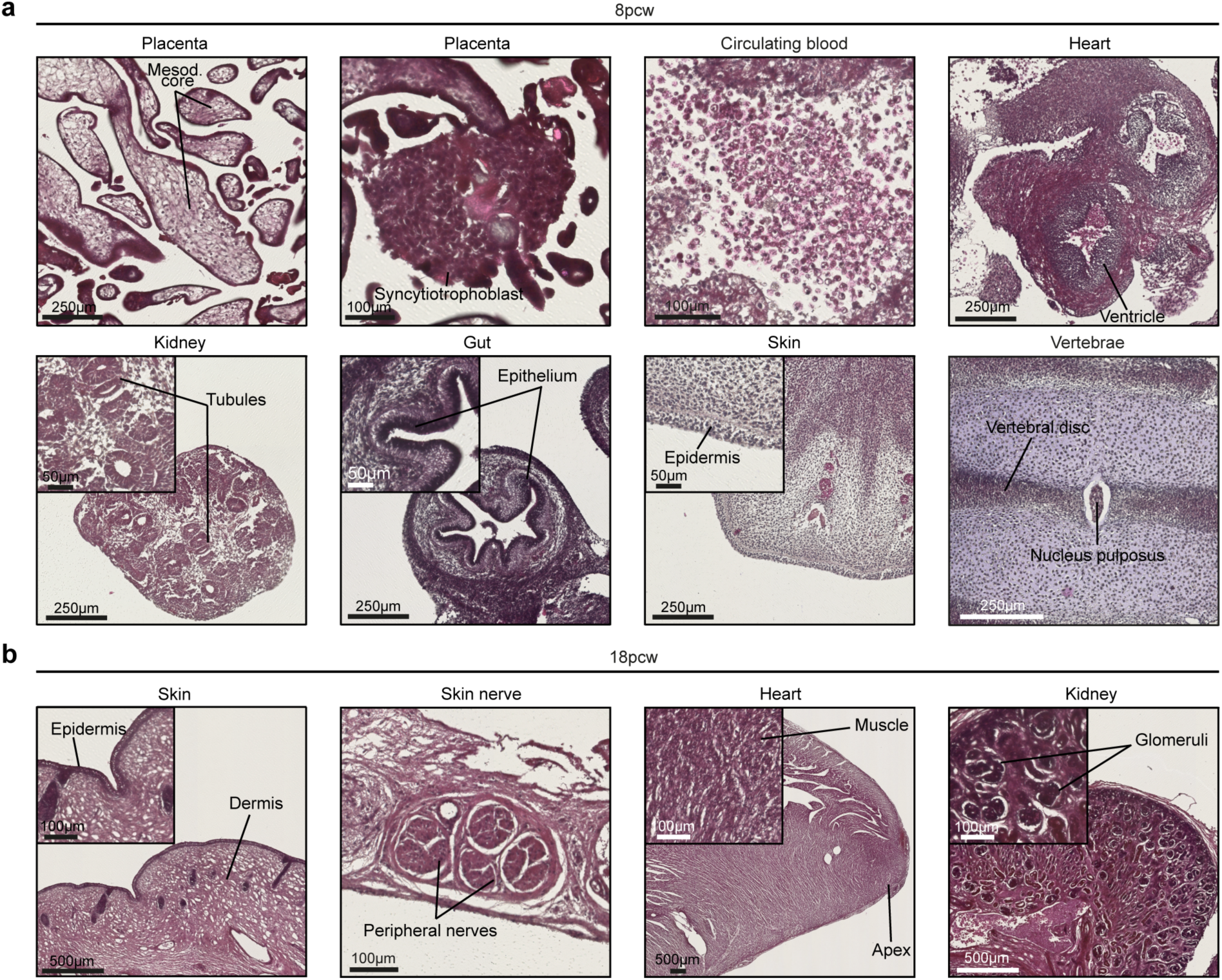
Laser-capture microdissected tissues. Representative histological slides of tissue structures with different developmental origins. Slides were stained with haematoxylin and eosin prior to microdissection. **a**, Structures microdissected from the 8 pcw sample included: mesodermal core and syncytiotrophoblast from the placenta, blood circulating in the heart, muscle from the heart, tubules from the kidney, epithelium from the gut, epidermis from the skin, vertebral disc from the vertebrae. **b**, Structures microdissected from the 18 pcw sample included: epidermis and peripheral nerves from the skin, muscle from the heart and glomeruli from the kidney.

**Extended Data Fig. 5.**
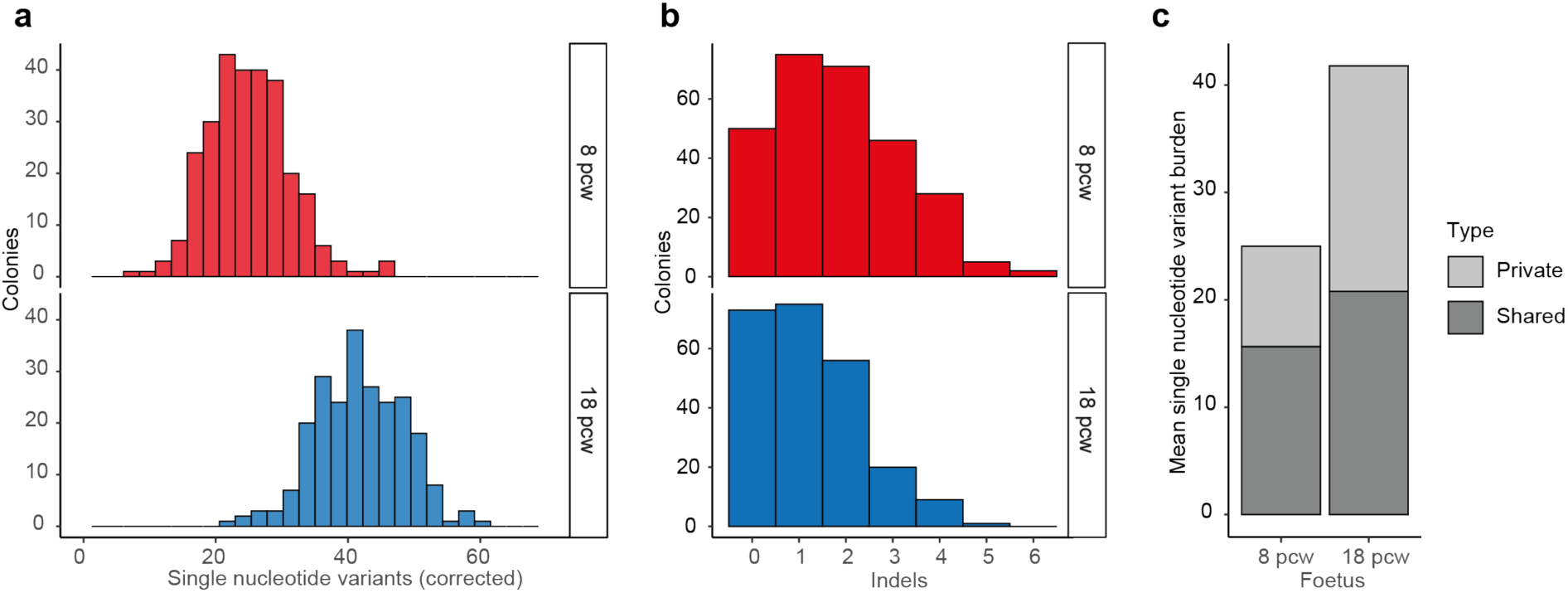
Mutation burden of single-cell derived colonies. Histograms showing: **a**, SNV burden per single-HSPC derived colony for the 8 pcw and 18 pcw foetuses, corrected for both the proportion of private mutations that are acquired in vitro, and for sample sensitivity. **b**, Indel burden per single-HSPC derived colony for the 8 pcw and 18 pcw foetuses, with no correction applied. **c**, Mean numbers of private and shared SNVs per colony for each foetus.

**Extended Data Fig. 6.**
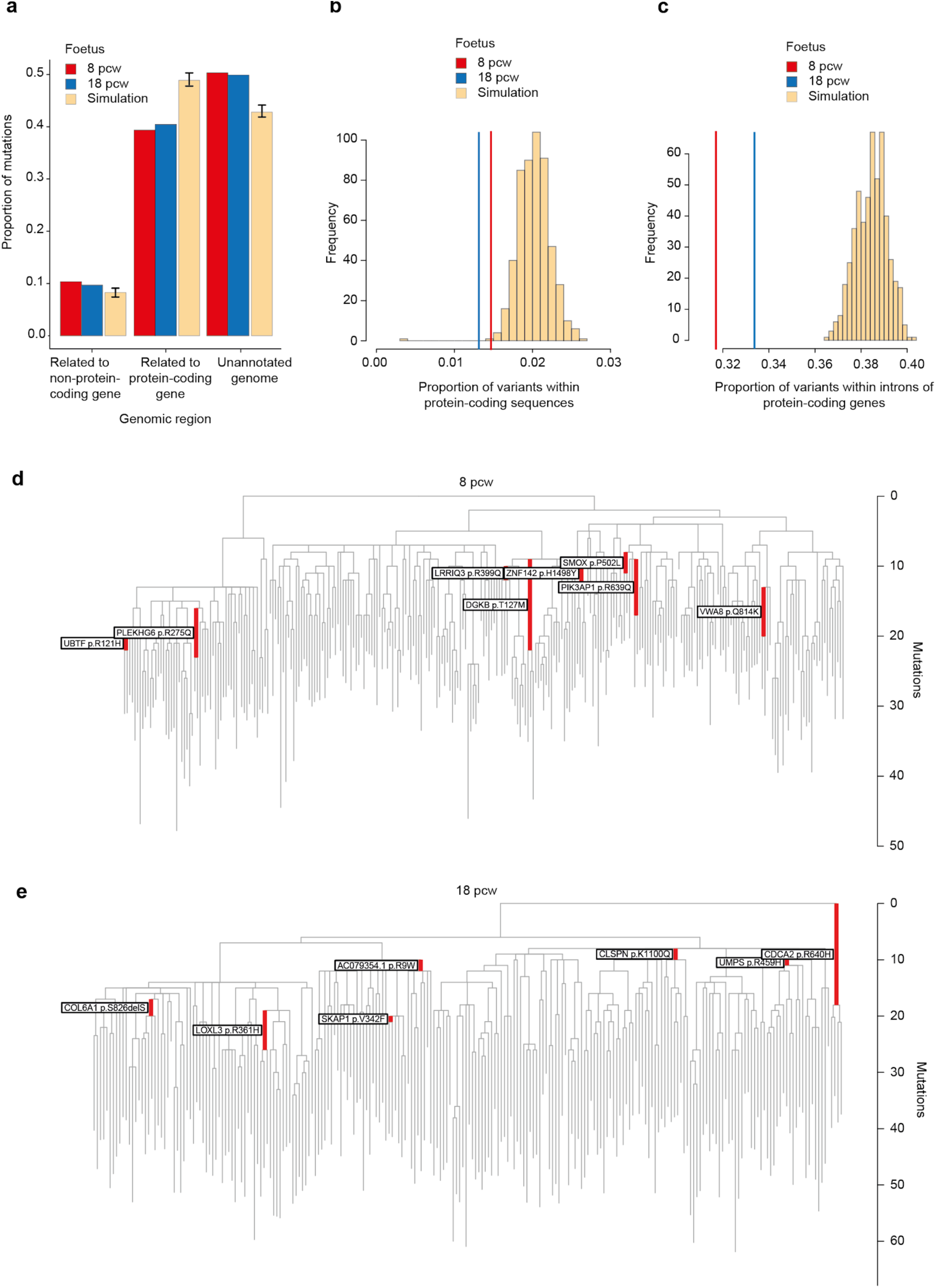
Genomic distribution of identified somatic mutations. **a**, Bar plot showing the proportion of variants mapping to different genomic features. **b**, Plot showing the proportion of variants within protein-coding sequences. The histograms show the distribution obtained by 500 simulations of random acquisition of mutations across the genome. **c**, Plot showing the proportion of variants within introns of protein-coding genes. The histograms show the distribution obtained by 500 simulations of random acquisition of mutations across the genome. **d**, Mutations in protein-coding sequences in the 18 pcw foetus, mapped on the branch of acquisition. Only those found in ≥2 colonies are shown. **e**, Mutations in protein-coding sequences in the 8 pcw foetus, mapped on the branch of acquisition. Only those found in ≥2 colonies are shown.

**Extended Data Fig. 7.**
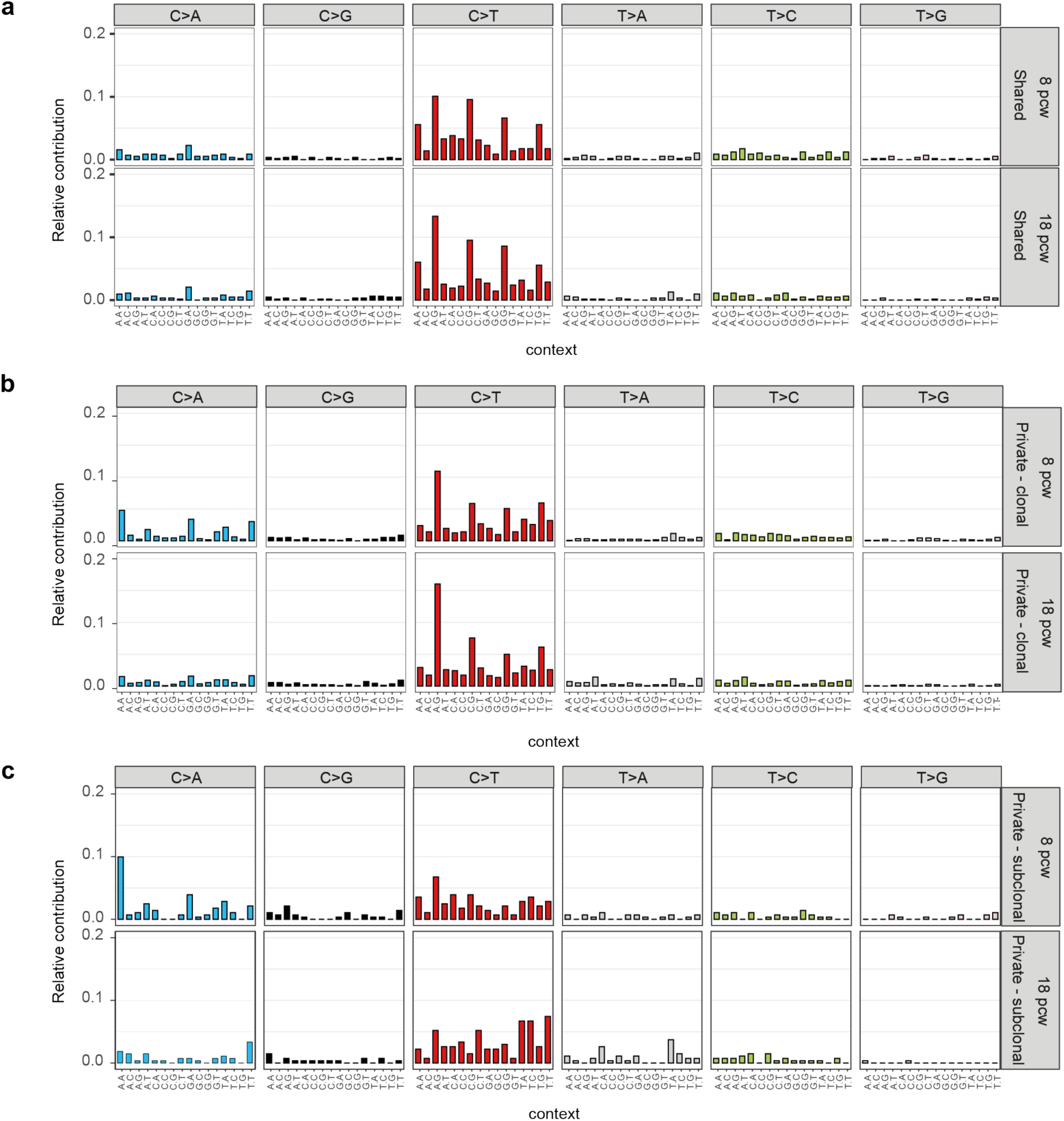
Mutational signatures. Mutational signatures incorporating the trinucleotide context of **a**, Shared mutations. **b**, Private mutations assigned to the clonal peak in the binomial mixture model, i.e. likely in vivo acquired mutations. **c**, Private mutations assigned to sub-clonal peaks in the binomial mixture, i.e. likely in vitro acquired mutations.

**Extended Data Fig. 8.**
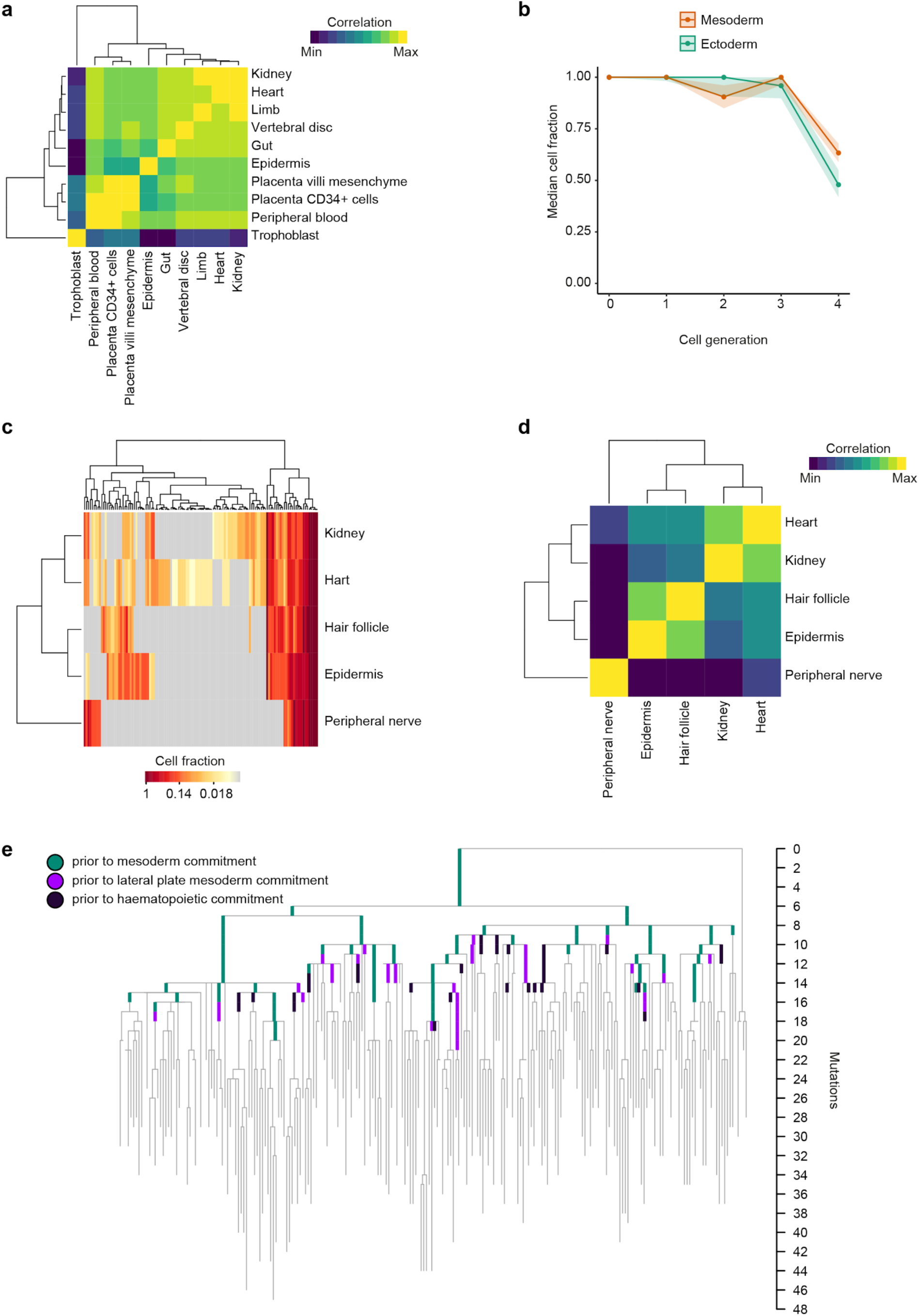
**a**, Heatmap representation of the targeted sequencing data from the 8 pcw foetus. Each column and each row represents a single tissue. The colour shows the level of correlation. **b**, Line plot showing lineage loss of microdissected tissues in the 18 pcw foetus at different times (represented as cell generations from the zygote). Shaded areas represent confidence intervals. **c**, Heatmap representation of the targeted sequencing data from the 18 pcw foetus. Each column represents an individual mutation, and each row a single tissue. The colour shows the VAF of the mutation in that tissue with grey indicating that it was not detected. **d**, Heatmap representation of the targeted sequencing data from the 18 pcw foetus. Each column and each row represents a single tissue. The colour shows the level of correlation. **e**, Phylogenetic tree showing the clonal relationships of 18 pcw HSPCs. Mutations identified in HSPCs by WGS that were also detected in non-haematopoietic tissues are coloured on the tree according to the earliest diverging tissue in which the mutation was reliably detected. Branch lengths are proportional to the number of mutations accumulated. The tree only contains those SNVs included in the bait set for targeted sequencing. ICM = inner cell mass. In all heatmaps of this figure the tissues are clustered using soft cosine similarity (see methods).

**Extended Data Fig. 9.**
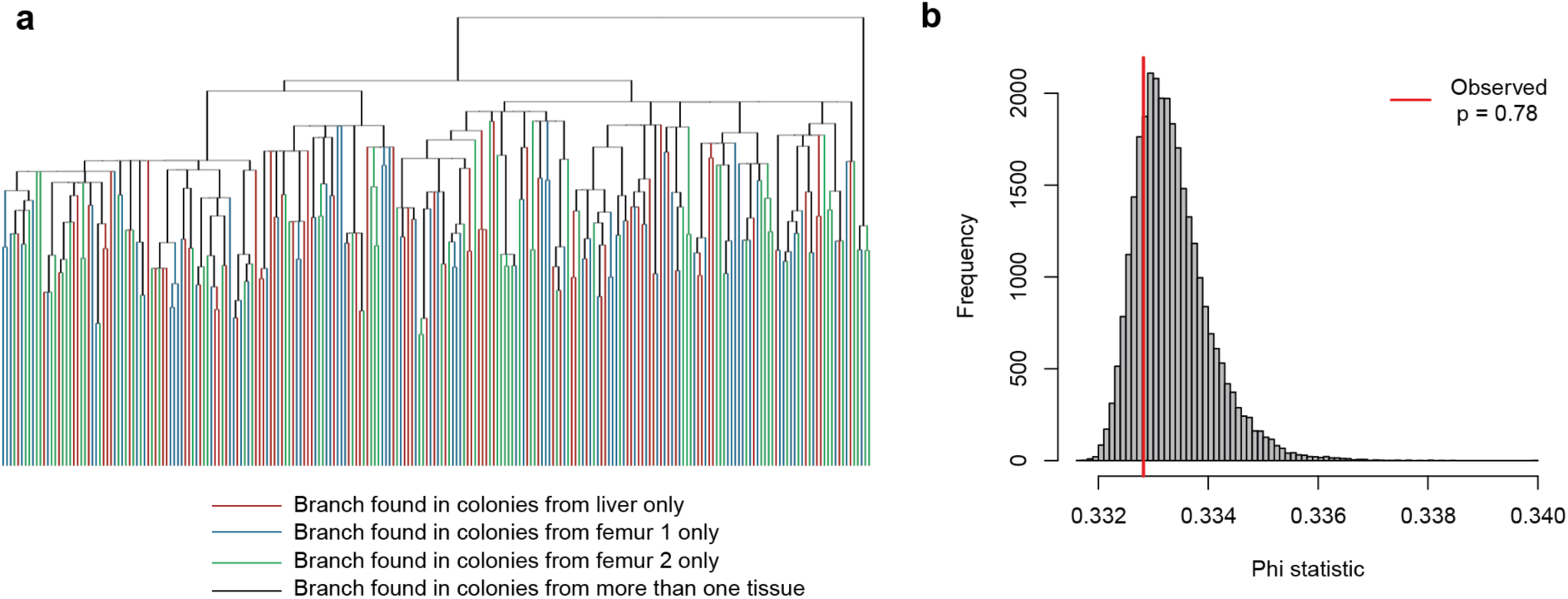
High level of intermixing of 18 pcw HSPCs among fetal liver and two sites of bone marrow. **a**, Ultrametric phylogenetic tree of 18 pcw HSPCs with branches coloured by the tissue from which HSPCs bearing the mutations were isolated. Black branches indicate that cells were found in more than one tissue, coloured branches indicate cells were unique to one specific tissue. **b**, Analysis of molecular variance used to formally test for clustering on the phylogeny of HSPCs isolated from the same tissue. The histogram shows the null distribution used to detect clustering. Distributions were obtained by randomly permuting which cells were assigned to which tissue. The observed value of the phi statistic is shown as a red line, with the p-value indicating no statistically significant clustering by tissue.

**Extended Data Table 1.**
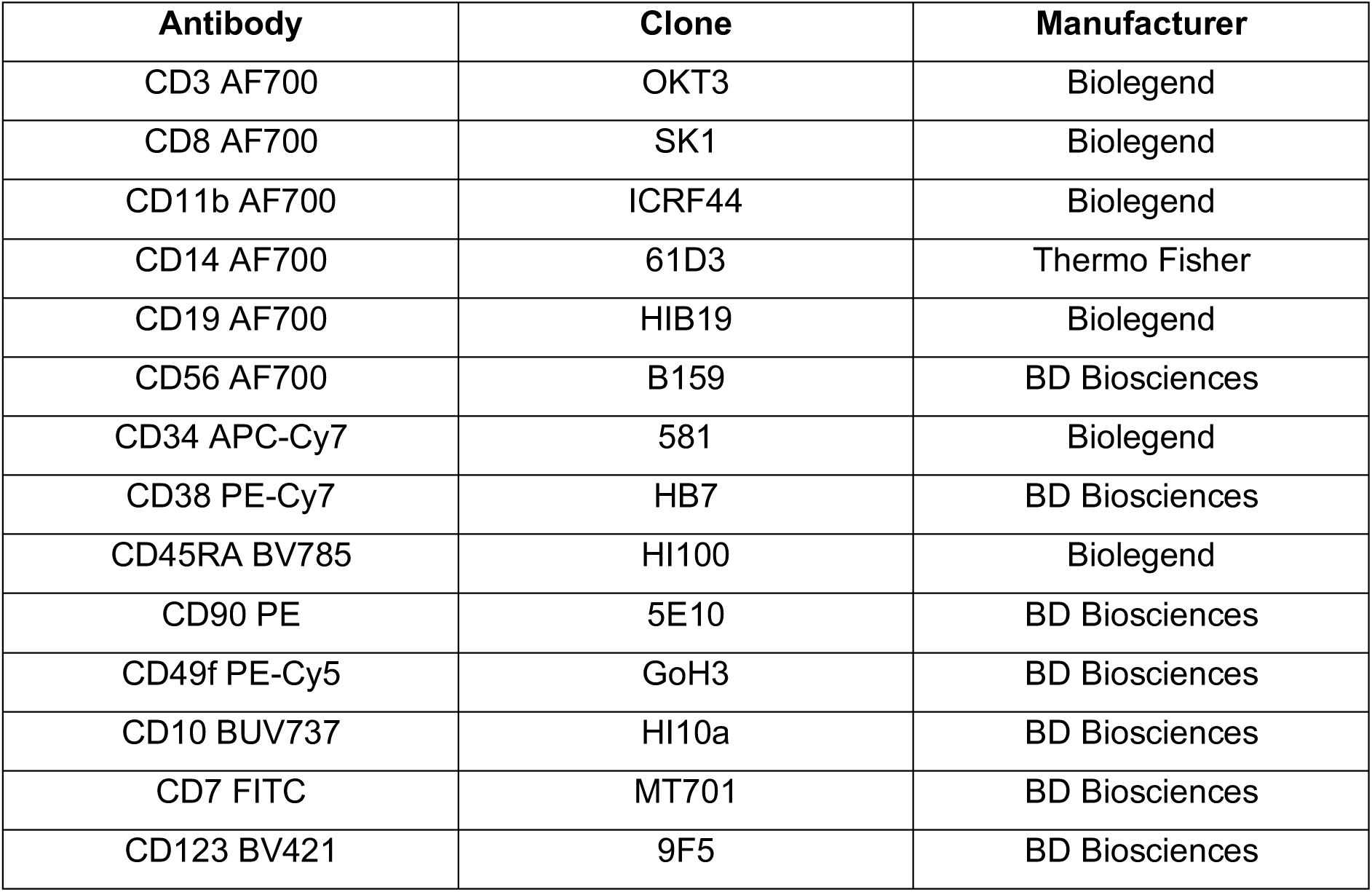
Antibodies list. Antibodies used for the study, including clone and manufacturer.

**Extended Data Table 2.** Laser capture microdissection biopsies. The number of biopsies undergoing targeted sequencing from each tissue for each foetus. The mean aggregated depth is the mean depth across all sites included in the custom capture panel after aggregating reads across biopsies from a given tissue.

| **Tissue**<br>**(8 pcw fetus)** | **Number of biopsies**<br>**undergoing sequencing** | **Mean aggregate depth**<br>**across biopsies (x)** |
| --- | --- | --- |
| Epidermis | 6 | 314 |
| Gut epithelium | 12 | 841 |
| Heart | 9 | 449 |
| Intervertebral disc | 4 | 184 |
| Kidney | 10 | 662 |
| Limb | 6 | 390 |
| Mesenchymal core of placenta | 5 | 152 |
| Peripheral blood | 5 | 268 |
| Trophoblast | 10 | 652 |
| Bulk FACS-sorted placental HSPCs | 1 | 78 |

**Extended Data Table 2 | Laser capture microdissection biopsies.**
| **Tissue**<br>**(18 pcw fetus)** | **Number of biopsies**<br>**undergoing sequencing** | **Mean aggregate depth**<br>**across biopsies (x)** |
| --- | --- | --- |
| Epidermis | 10 | 551 |
| Hair follicle | 4 | 239 |
| Heart | 10 | 1246 |
| Kidney | 13 | 701 |
| Peripheral nerve | 1 | 32 |

